# A pro-metastatic tRNA fragment drives Nucleolin oligomerization and stabilization of bound metabolic mRNAs

**DOI:** 10.1101/2021.04.26.441477

**Authors:** Xuhang Liu, Hanan Alwaseem, Henrik Molina, Bernardo Tavora, Sohail F. Tavazoie

## Abstract

Stress-induced cleavage of transfer RNAs (tRNAs) into tRNA-derived fragments (tRFs) occurs across organisms from yeast to human, yet its mechanistic bases and pathological consequences remain poorly defined. By performing genome-wide small RNA profiling, we detected increased abundance of a Cysteine tRNA fragment (5’-tRF^Cys^) during breast cancer metastatic progression. 5’’-tRF^Cys^ is required for efficient breast cancer metastatic lung colonization and metastatic cell survival. We identified Nucleolin as the direct binding partner of 5’-tRF^Cys^. 5’-tRF^Cys^ binding enhanced the stability of Nucleolin’s associated pro-metastatic transcripts encoding metabolic enzymes Mthfd1l and Pafah1b1. 5’-tRF^Cys^ stabilized these transcripts by promoting Nucleolin oligomerization and the assembly of Nucleolin and its bound transcripts into a higher-order ribonucleoprotein complex. Our findings reveal that a tRF can promote oligomerization of an RNA binding protein into a stabilizing ribonucleoprotein complex containing specific target transcripts, thereby driving specific metabolic pathways underlying cancer progression.

## INTRODUCTION

Transfer RNA-derived RNA fragments (tRFs) are a novel class of small non-coding RNAs that are cleaved from mature or precursor transfer RNAs (tRNAs). TRFs are classified into distinct groups based on their sites of origin within tRNAs. 5’- and 3’-tRNA halves arise from cleavage within the anticodon loop by Angiogenin, RNase L or RNase 1 (Donovan et al., 2017; Fu et al., 2009; Nechooshtan et al., 2020; Yamasaki et al., 2009). Additionally, 5’- and 3’-tRF-18 are 18-nt fragments generated by Dicer (Babiarz et al., 2008; Cole et al., 2009), while 3’U tRFs are cleaved by RNase Z from the 3’ end of precursor tRNAs (Lee et al., 2009). TRFs are important regulators of a number of molecular and cellular processes (Oberbauer and Schaefer, 2018; Schimmel, 2018). For example, a subset of hormone-stimulated 5’-tRFs are required for proliferation in breast and prostate cancer cells (Honda et al., 2015). 5’-tRFs influence transgenerational inheritance of metabolism in mice (Chen et al., 2016; Sharma et al., 2016). 3’-tRF^Lys^ inhibits transposition of retroviruses by serving as a decoy for tRNA^Lys^ (Schorn et al., 2017). 3’-tRF^Ala^ enhances translation by upregulating the expression of a key ribosomal protein gene (Kim et al., 2017). In response to hypoxia, a subset of tRFs containing a common motif were found to become upregulated in cancer cells and bind the RNA-binding protein Y-Box Binding Protein 1 (YBX1)—competitively inhibiting YBX1 binding to its growth-promoting target transcripts and consequently suppressing cancer progression (Goodarzi et al., 2015). Despite their demonstrated roles in a variety of cellular processes, the molecular mechanisms underlying tRF function remain poorly defined.

Nucleolin is an evolutionarily conserved RNA binding protein that plays crucial roles in multiple molecular processes (Mongelard and Bouvet, 2007). Nucleolin is essential for ribosomal biogenesis, ribosomal RNA (rRNA) transcription and processing, and assembly of ribosomes (Ginisty et al., 1998; Ginisty et al., 1999; Serin et al., 1996). Nucleolin also regulates multiple steps in messenger RNA (mRNA) processing, including splicing, stabilization, nucleus-cytoplasmic transport and translation (Abdelmohsen and Gorospe, 2012). Recent studies have further implicated Nucleolin in the synthesis of microRNAs (miRNAs) (Pichiorri et al., 2013; Pickering et al., 2011) and self-renewal of embryonic stem cells (Percharde et al., 2018). In keeping with its pleiotropic roles in post-transcriptional control, Nucleolin is essential for the proliferation and survival of mammalian cells (Ugrinova et al., 2007). Despite decades of research, the full spectrum of RNAs directly bound to Nucleolin has remained unclear.

Post-transcriptional regulation, which is carried out by RNA binding proteins (RBPs) and non-coding RNAs, is a critical step in gene expression (Corley et al., 2020). An emerging paradigm for the proper functioning of RBPs is their oligomerization. Formation of higher-order RNA-RBP complexes plays a critical role for pattern recognition receptors RIG-I and MDA5 to trigger immune response against foreign nucleic acids (Jiang et al., 2012; Patel et al., 2013; Peisley et al., 2011; Wu et al., 2013), and for splicing factor Rbfox and multivalent hnRNPs to regulate alternative splicing (Gueroussov et al., 2017; Ying et al., 2017). Oligomeric RBPs can also serve as the building blocks for the assembly of filaments (Peisley et al., 2011; Peisley et al., 2014; Peisley et al., 2013; Wu et al., 2013) or membrane-less RNA/ribonucleoprotein granules, which are large assemblies formed by RNA-RNA, RNA-RBP and RBP-RBP interaction (Alberti and Hyman, 2021; Lyon et al., 2021; Weber and Brangwynne, 2012). It was proposed that long RNAs promote RBP oligomerization through intermolecular RNA-RNA interactions and by serving as a scaffold for multivalent RBPs (Van Treeck and Parker, 2018). Given the vast number of small RNAs in the genome, it is currently unknown what role, if any, small RNAs play in the formation of higher-order RNA-RBP complexes.

During cancer progression, there is a selection for cancer cells that overcome multiple restrictive physiological barriers at the primary and metastatic microenvironments. These barriers are bypassed when rare cells modulate gene expression programs to enable multiple adaptive phenotypes (Hanahan and Weinberg, 2011). A critical mechanism by which such gene expression states are achieved is through alterations in the expression levels of specific small RNAs, which enables pro-metastatic, post-transcriptional gene expression responses (Loo et al., 2015; Mendell and Olson, 2012; Nabet et al., 2017; Pencheva et al., 2012; Truitt and Ruggero, 2017). We hypothesized that certain tRFs may become induced during metastatic progression. We reasoned that by studying how such tRFs shape gene expression programs, we might uncover insights into the mechanisms of action of these molecules. Herein, we report that a tRNA fragment, 5’-tRF^Cys^, derived from the 5’ half of Cysteine tRNA is upregulated in multiple highly metastatic breast cancer cell lines and primary human breast tumors. 5’-tRF^Cys^ is required for breast cancer metastatic colonization and survival. We find that this fragment directly binds to Nucleolin, enhances the interaction of Nucleolin with specific target transcripts and promotes the formation of an oligomeric Nucleolin/transcript complex—collectively stabilizing the Nucleolin-bound transcripts. 5’-tRF^Cys^ and Nucleolin promotes breast cancer metastasis by stabilizing transcripts encoding Pafah1b1 and Mthfd1l, key metabolic enzymes. Our results uncover a 5’-tRF^Cys^/Nucleolin axis that drives breast cancer metastatic progression and demonstrate that a tRF can enhance oligomerization and post-transcriptional function of an RNA binding protein.

## RESULTS

### 5’-tRFCys is upregulated during breast cancer progression and metastasis

To search for tRFs that may regulate breast cancer metastasis, we performed next-generation sequencing of small RNAs isolated from three isogenic mouse breast cancer cell lines with differing metastatic capacities — highly metastatic 4T1, poorly metastatic 4TO7, and non-metastatic 67NR cells (Aslakson and Miller, 1992; Dexter et al., 1978). Hierarchical cluster analysis based on sample-to-sample distance revealed that the expression levels of small RNAs were sufficiently informative to classify the samples into their respective highly, poorly and non-metastatic groups (Figure S1A). Among different classes of tRFs, the most abundant species were 5’-tRNA halves (Figure S1B). We found 5’-tRF^Cys^, derived from the 5’ half of multiple tRNA^Cys^GCA isodecoders, to be one of the most upregulated 5’ tRNA halves in highly metastatic 4T1 cells relative to isogenic poorly metastatic 4TO7 cells (Figure 1A). Quantitative reverse transcription PCR (RT-qPCR) using a 5’-tRF^Cys^ specific stem-loop primer independently confirmed that expression of 5’-tRF^Cys^ was significantly elevated in 4T1 relative to 4TO7 or 67NR cells (Figure 1B). A similar increase in 5’-tRF^Cys^ expression was observed in an independent *in vivo* selected EO771-LM3 highly metastatic cell line relative to its parental EO771-Par poorly metastatic isogenic line (Figure 1C). These results reveal that 5’-tRF^Cys^ is upregulated in breast cancer cells of high metastatic potential.

**Figure 1.**
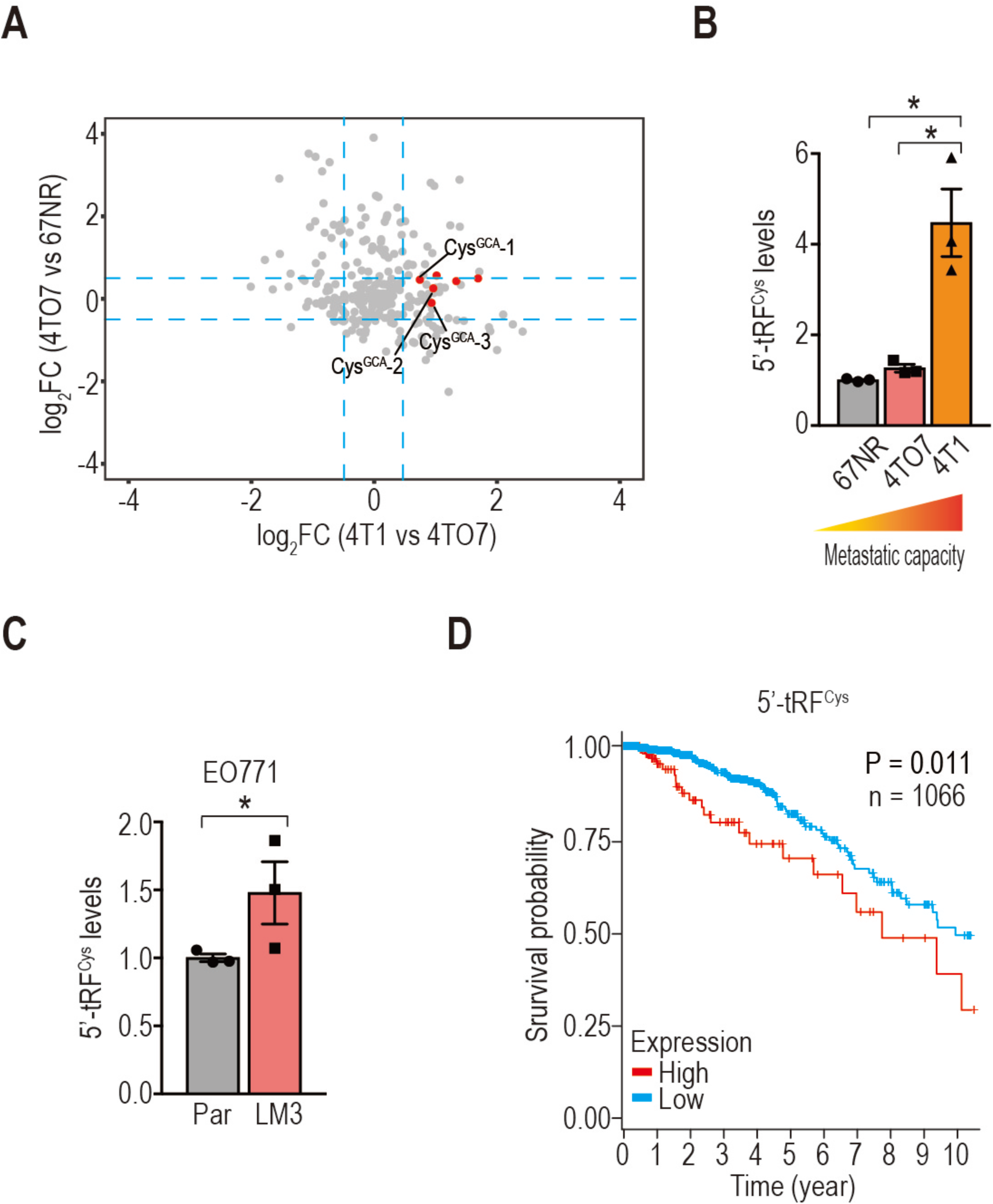
5’-tRF^Cys^ is upregulated during breast cancer progression and metastasis. A. A scatter plot depicting log_2_FoldChange (log_2_FC) for tRNA fragments isolated from highly metastatic 4T1, poorly metastatic 4TO7, and non-metastatic 67NR cells. 5’-tRNA halves that were significantly upregulated only in 4T1 but not 4TO7 or 67NR cells are marked in red. The blue dashed lines denote −0.5 and 0.5, respectively. B, C. Quantification of 5’-tRF^Cys^ levels by RT-qPCR from mouse breast cancer cells with differing metastatic capacities (N=3). All data hereafter are represented as mean ± s.e.m. and data points represent biological replicates. Representative results from at least two independent experiments are shown, except mouse experiments, most of which were performed once with cohorts of mice. Statistical significance is determined by one-tailed t-test with Welch’s correction. *, p<0.05. D. Kaplan-Meier curve depicting survival probability of breast cancer patients (n=1066) in the TCGA cohort stratified by 5’-tRF^Cys^ expression levels. Statistical significance is determined by the Mantel–Cox log-rank test.

Analysis of small RNA sequencing data from the Cancer Genome Atlas (TCGA) (Cancer Genome Atlas, 2012; Pliatsika et al., 2018) revealed that high expression of 5’-tRF^Cys^ correlated with poor survival of breast cancer patients (Figure 1D; P=0.011; n=1066). Additionally, the expression of 5’-tRF^Cys^ is significantly elevated in breast cancer samples relative to matched normal breast tissues in the TCGA cohort (Figure S1C; P=1.82e-11; n=101). Together, these results reveal that 5’-tRF^Cys^ expression is increased during breast cancer progression.

### 5’-tRF^Cys^ is required for efficient metastatic lung colonization of breast cancer cells

To test whether 5’-tRF^Cys^ plays a functional role in breast cancer metastasis, we sought to inhibit the activity of 5’-tRF^Cys^ using two distinct antisense locked nucleic acid (LNA) oligonucleotides. Such LNAs inhibit small RNA function by binding target mRNAs and impairing their interaction with target RNAs or proteins. In some cases, such LNAs can functionally deplete target RNAs (Kim et al., 2017). Indeed, we observed such a depletion effect upon use of one 5’-tRF^Cys^-targeting LNA (Figure S2A). Transfection of either antisense LNA oligonucleotide into two highly metastatic mouse breast cancer cell lines (4T1 and EO771-LM3) significantly reduced metastasis in tail-vein colonization assays (Figures 2A, 2B and S2B). Additionally, inhibiting 5’-tRF^Cys^ by transducing an antisense tough decoy molecule (Haraguchi et al., 2009; Xie et al., 2012) in the highly metastatic MDA-MB-231-LM2 human breast cancer cell line and a human patient-derived xenograft organoid (PDXO) line significantly suppressed metastatic colonization in immunocompromised NOD-scid gamma (NSG) mice (Figures S2C and S2D). These results reveal that 5’-tRF^Cys^ plays a critical role in breast cancer metastatic colonization independent of an intact immune system.

**Figure 2.**
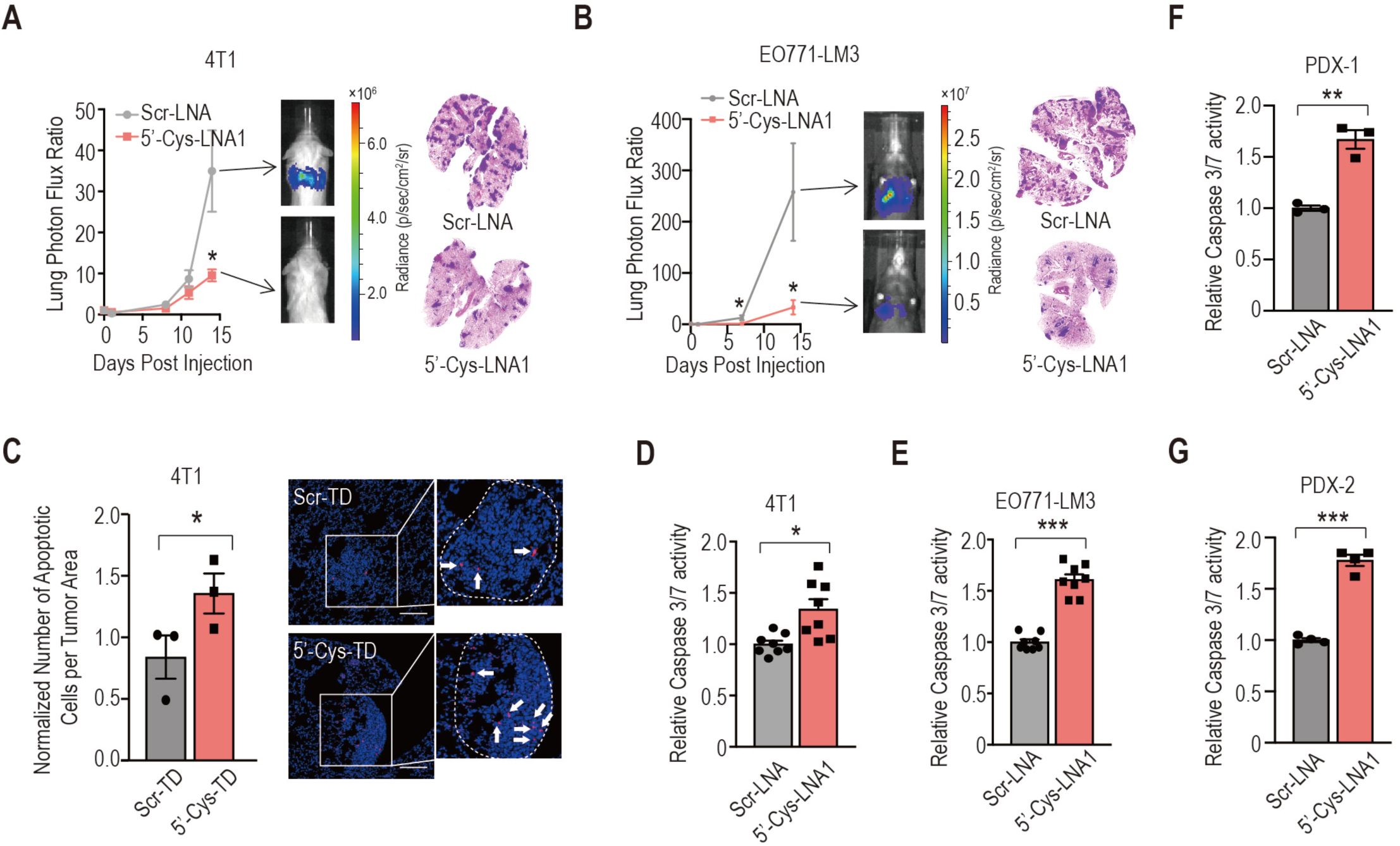
5’-tRF^Cys^ promotes breast cancer metastasis and enhances breast cancer cell survival. A, B. Bioluminescence imaging plots of metastatic lung colonization by 4T1 (A) and EO771-LM3 (B) cells transfected with a scrambled control (Scr-LNA) or a 5’-tRF^Cys^ antisense LNA oligo (5’-Cys-LNA1). Representative bioluminescence images and H&E staining of lung sections for each cohort are shown (N=5-7). C. Left, quantification of cleaved Caspase 3 positive cells in metastatic lung nodules normalized by areas of lung nodules. Mice were injected with 4T1 cells transduced with a 5’-tRF^Cys^ antisense (5’-Cys-TD) or a scrambled control tough decoy (Scr-TD) (N=3). Each data point represents the average of at least 10 different image measurements from one mouse lung section. Right, representative confocal images of anti-cleaved Caspase 3 staining from mouse lung sections. The dashed line delineates metastatic nodules. Scale bar: 10 um. D, E, F, G. Quantification of Caspase3/7 activity in 4T1 (D), EO771-LM3 (E) cells or two human breast cancer patient derived xenograft organoid lines (F&G) upon suppression of 5’-tRF^Cys^ (N=3-8). Statistical significance in mouse (A and B) and cell biology (C-G) experiments is determined by Mann-Whitney test and one-tailed t-test with Welch’s correction, respectively. *, p<0.05; **, p<0.01; ***, p<0.001.

We next determined the cellular mechanism by which 5’-tRF^Cys^ promotes metastasis. Immunostaining of metastatic lung sections revealed no significant difference in the abundance of Ki-67 or Endomucin positive cells (Figures S2E and S2F) upon 5’-tRF^Cys^ inhibition, suggesting that neither proliferation nor angiogenesis was affected. In contrast, we observed a significant increase in the number of cleaved Caspase-3 positive foci in metastatic lung nodules upon 5’-tRF^Cys^ inhibition (Figure 2C). Consistent with this, suppressing 5’-tRF^Cys^ significantly increased Caspase3/7 activity in 4T1 and EO771-LM3 metastatic cells as well as two human PDXO lines *in vitro* (Figures 2D-2G). These findings reveal that 5’-tRF^Cys^ regulates metastatic colonization by promoting survival of breast cancer cells.

### Nucleolin directly binds 5’-tRF^Cys^

To search for the mechanism underlying 5’-tRF^Cys^’s critical role in metastasis, we attempted to identify protein(s) that directly interact with 5’-tRF^Cys^ using a biochemical approach, since small non-coding RNAs generally mediate their effects through interactions with RNA-binding proteins (RBPs). As such, we irradiated cells with UV to crosslink proteins to RNA substrates and used a biotinylated 5’-tRF^Cys^ antisense oligonucleotide to pull down 5’-tRF^Cys^-interacting partners from 4T1 cell lysates. Mass spectrometry (MS) identified Nucleolin as the most enriched protein in the 5’-tRF^Cys^ antisense pulldown relative to the antisense control (Figure 3A). In reciprocal pulldown experiments using an antibody against Nucleolin, we observed a ∼500-fold enrichment of 5’-tRF^Cys^ from Nucleolin immunoprecipitates (IP) relative to the control IP (Figure 3B). These experiments demonstrate that that Nucleolin is a binding partner of 5’-tRF^Cys^.

**Figure 3.**
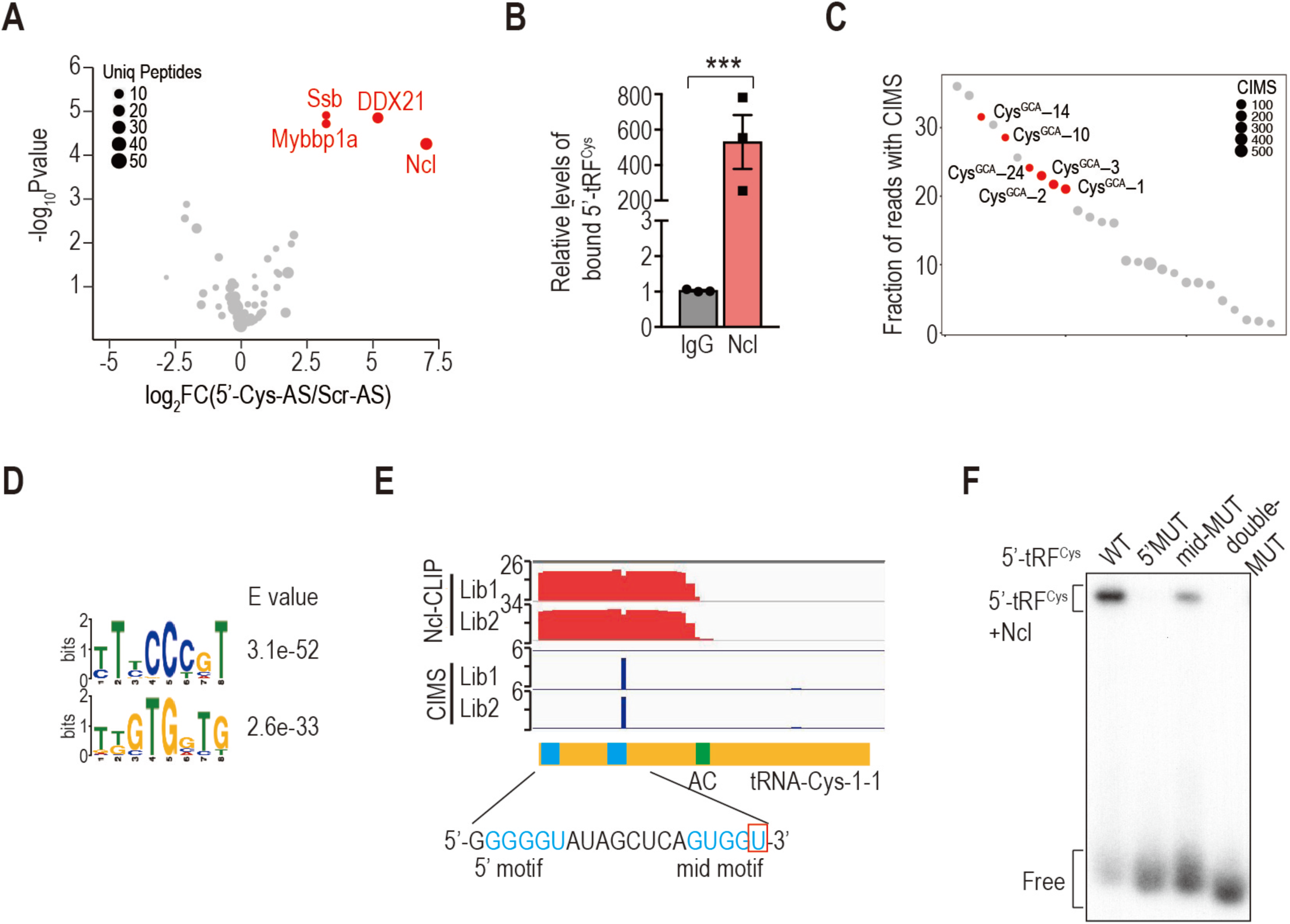
Nucleolin is a direct binding partner of 5’-tRF^Cys^. A. Volcano plot depicting log_2_FC values versus –log_10_Pvalue in protein abundance quantified from pulldowns conducted with a biotinylated 5’-tRF^Cys^ antisense (5’-Cys-AS) in comparison to a biotinylated scrambled control oligo (Scr-AS) in 4T1 cells. The most enriched proteins in the antisense group are marked in red. B. Quantification of the amount of 5’-tRF^Cys^ pulled down by a Nucleolin or an IgG control antibody using Taqman qRT-PCR assays. Statistical significance was determined by one-tail t-tests with Welch’s correction. ***, p<0.001. C. Dot plot depicting 5’-tRNA halves with the highest fraction of crosslinking induced modification sites (CIMS) in the non-RNase-treated Nucleolin CLIP libraries. Reads derived from the tRNA^Cys^ loci are marked in red. D. The two most enriched motifs identified using Nucleolin-bound CLIP tags with CIMS sites in non-RNase-treated Nucleolin CLIP libraries. E. Genome browser view of Nucleolin-CLIP tags and CIMS sites in a representative tRNA^Cys^ locus from two non-RNase treated libraries. The blue and green boxes denote the two G-rich motifs and the anticodon (AC), respectively. The red rectangle marks the CIMS site. The Y axis in the CLIP tag track represents reads per million (RPM) while that in the CIMS track represents the actual number of CIMS sites. F. Electrophoresis mobility shift assay (EMSA) in the presence of an EDTA-containing buffer using purified Nucleolin protein and 5’ radiolabeled wild-type (WT) or mutant (MUT) 5’-tRF^Cys^ containing mutations in the 5’ (5’MUT), middle (mid-MUT) or both (double-MUT) G-rich motifs.

To test whether Nucleolin directly binds 5’-tRF^Cys^, we performed high-throughput sequencing of RNA isolated by crosslinking immunoprecipitation (HITS-CLIP) (Moore et al., 2014) in 4T1 cells. Notably, unlike canonical HITS-CLIP, no exogenous ribonuclease (RNase) was used in this experiment in order to preserve the natural ends of small RNAs. Prior studies have shown that the fraction of reads with crosslinking induced modification sites (CIMS) correlates with the binding affinity of an RBP for its substrate RNA (Zhang and Darnell, 2011). Bioinformatic analyses revealed that 5’-tRNA halves were the most enriched type of tRFs bound to Nucleolin (Figure S3A). Importantly, several of the most enriched 5’-tRNA halves were derived from tRNA^Cys^ loci (Figure 3C). These results reveal a high binding affinity for Nucleolin towards 5’-tRF^Cys^, implicating Nucleolin as a direct binding partner of 5’-tRF^Cys^.

Motif analysis of sequences surrounding CIMS sites revealed the enrichment of two motifs (Figure 3D). The first C-rich motif is highly similar to the Nucleolin recognition element, which was previously identified by *in vitro* biochemical assays and plays a critical role in mediating Nucleolin’s binding to the 5’ external transcribed spacers (ETS) of pre-45S rRNA transcripts (Ghisolfi-Nieto et al., 1996). The second G-rich motif has not been previously reported. 5’-tRF^Cys^ contains a CIMS site right next to this motif (Figure 3E), suggesting that this motif may mediate Nucleolin binding to 5’-tRF^Cys^. Interestingly, we noted that there was another highly similar G-rich motif at the 5’ end of 5’-tRF^Cys^. To test whether these motifs are required for Nucleolin binding to 5’-tRF^Cys^, we mutated each individually or in combination. Electrophoretic mobility shift assays (EMSA) with purified Nucleolin protein (Figure S3B) revealed that mutating either G-rich motif of 5’-tRF^Cys^ dramatically reduced Nucleolin’s binding (Figure 3F). These findings reveal that two G-rich motifs of 5’-tRF^Cys^ are required for the interaction between Nucleolin and 5’-tRF^Cys^.

### 5’-tRFCys facilitates recruitment of Nucleolin to target transcripts

Given the well-established role for Nucleolin as an essential protein in ribosomal RNA (rRNA) biogenesis (Srivastava and Pollard, 1999), we next examined whether the binding of 5’-tRF^Cys^ to Nucleolin affects translation. We observed that neither rRNA synthesis nor global translation was impaired upon inhibition of 5’-tRF^Cys^ (Figures S4A and S4B).

Considering that Nucleolin has also been previously implicated in regulating mRNA stability and translation (Abdelmohsen et al., 2011; Sengupta et al., 2004; Takagi et al., 2005; Zaidi and Malter, 1995), we postulated that 5’-tRF^Cys^ may regulate Nucleolin binding to specific mRNAs. To test this hypothesis, we performed HITS-CLIP of Nucleolin in cells transfected with either a scrambled control or 5’-tRF^Cys^ antisense LNA oligonucleotide. Surprisingly, in addition to the well-established targets such as rRNAs and snoRNAs, we found that Nucleolin bound a wide spectrum of mRNA transcripts (Figure S4C), with the majority of binding peaks residing within 5’ untranslated regions (UTRs) (Figure S4D). Interestingly, Nucleolin binding to a subset of peaks was significantly reduced upon 5’-tRF^Cys^ inhibition (Figure 4A), suggesting that Nucleolin’s binding to a subset of transcripts was promoted by 5’-tRF^Cys^.

**Figure 4.**
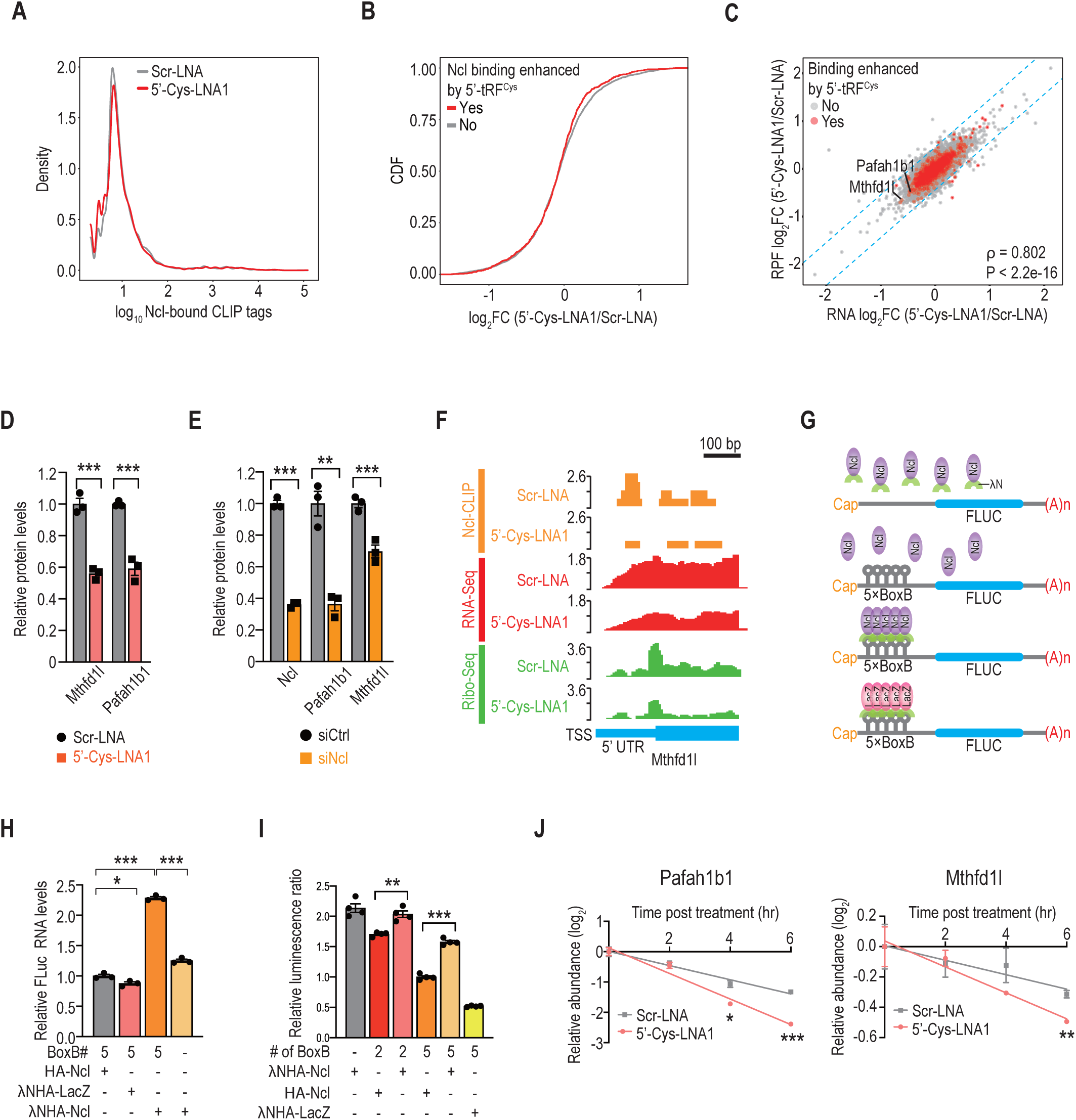
5’-tRF^Cys^ promotes Nucleolin binding to its target transcripts to enhance their stability. A. Density plot of the log_10_ CLIP tag counts in Nucleolin-bound peaks identified from cells transfected with the control LNA (Scr-LNA) or 5’-tRF^Cys^ antisense LNA (5’-Cys-LNA1) oligos. Statistical significance was determined by Kolmogorov–Smirnov test (P < 2.2e-16). B. Cumulative distribution function (CDF) plots of the log_2_FC in protein abundance between 5’-tRF^Cys^ suppressed and control cells for all transcripts stratified by whether their Nucleolin binding was enhanced by 5’-tRF^Cys^ (red) or not (grey). Statistical significance was determined by the Kolmogorov–Smirnov test (P = 0.037). C. Scatter plot comparing log_2_FC in the number of ribosome protected fragments (RPFs) and log_2_FC in the number of RNA-Seq reads between 5’-tRF^Cys^ suppressed (5’-Cys-LNA1) and control cells (Scr-LNA) for transcripts whose Nucleolin binding was enhanced by 5’-tRF^Cys^ (red) or not (grey). ρ, Spearman’s correlation coefficient. D, E. Quantification of 5’-tRF^Cys^ targets’ protein abundances upon inhibition of 5’-tRF^Cys^ (D) or depletion of Nucleolin (E). See also Figures S4H and S4I. F. Genome browser view of aligned Nucleolin (Ncl)-CLIP tags (orange), Ribo-Seq reads (green), and RNA-Seq reads (red) within the 5’ UTR of Mthfd1l. The Y-axis represents RPM. TSS, transcription start site. G. Schematic of the reporter assay wherein tethering a λN- and HA-tagged control protein (LacZ) or a λN- and/or HA-tagged Nucleolin protein to the firefly luciferase reporter with or without five BoxB sites. H, I. Quantification of reporter transcript abundance (H) and luciferase activity (I) by RT-qPCR (H) and dual luciferase assay (I), respectively. J. Quantification by RT-qPCR of the log_2_FC in Pafah1b1 and Mthfd1l transcript levels in control (Scr-LNA) or 5’-tRF^Cys^ suppressed cells (5’-Cys-LNA1) in the RNA stability assay. Statistical significance in D-E and H-J was determined by one-tail t-tests with Welch’s correction. *, p<0.05. **, p < 0.01. ***, p<0.001.

To examine the impact of depleting 5’-tRF^Cys^ on its target transcripts, we quantified proteome changes by mass spectrometry, focusing on genes whose transcripts exhibited enhanced 5’-tRF^Cys^-mediated Nucleolin binding. The protein levels of a subset of these genes were significantly reduced upon 5’-tRF^Cys^ inhibition (Figure 4B). These results suggest that 5’-tRF^Cys^ promotes Nucleolin binding to 5’-UTRs of target transcripts, thereby enhancing gene expression. In line with this hypothesis, we also found that Nucleolin-bound transcripts were significantly downregulated upon depletion of Nucleolin (Figure S4E), suggesting that Nucleolin binding stabilized its bound target transcripts.

To identify targets regulated by 5’-tRF^Cys^/ Nucleolin binding, we examined transcriptome and translatome changes upon 5’-tRF^Cys^ inhibition with two distinct 5’-tRF^Cys^ antisense LNAs by RNA Sequencing (RNA-Seq) and Ribosome Profiling (Ribo-Seq) respectively. Inhibition of 5’-tRF^Cys^ by the two distinct 5’-tRF^Cys^ antisense LNAs led to highly similar gene expression changes (Figure S4F). We found that both protein abundance and the abundance of ribosome protected fragments correlated significantly with their transcript levels (Figures 4C and S4G), suggesting that most Nucleolin-bound transcripts were regulated at the post-transcriptional rather than translational level. By looking for genes whose Nucleolin binding was enhanced by 5’-tRF^Cys^ and whose expression levels were downregulated in 5’-tRF^Cys^ inhibited cells, we identified two of the most downregulated Nucleolin-bound transcripts—Pafah1b1 and Mthfd1l— as candidate downstream effectors (Figure 4C). Platelet activating factor acetyl hydrolase 1 beta 1 (Pafah1b1) is a regulatory subunit of platelet activating factor acetyl hydrolase 1 beta (Karasawa and Inoue, 2015), whereas Methylenetetrahydrofolate Dehydrogenase 1 Like (Mthfd1l) is a metabolic enzyme that plays a crucial role in one-carbon metabolism (Ducker and Rabinowitz, 2017; Yang and Vousden, 2016) (Figure S6F). Importantly, both genes were found to promote progression of multiple cancer types in prior studies (Agarwal et al., 2019; Lee et al., 2017; Lo et al., 2012; Locasale, 2013; Selcuklu et al., 2012; Zimdahl et al., 2014). Western blotting confirmed that Pafah1b1 and Mthfd1l became downregulated upon inhibition of 5’-tRF^Cys^ (Figures 4D and S4H) or depletion of Nucleolin (Figures 4E and S4I).

Consistent with results from the aforementioned genome-wide analyses, Nucleolin predominantly bound Pafah1b1 and Mthfd1l in the 5’ UTRs of these transcripts in a 5’-tRF^Cys^-dependent manner (Figure 4F and S4J). RNA Sequencing revealed that both target transcripts became reduced upon 5’-tRF^Cys^ inhibition while Ribo-Seq analyses revealed a similar reduction in the number of ribosome-protected fragments (Figure 4F and S4J), suggesting that 5’-tRF^Cys^-mediated regulation of these two target candidates occurs at the post-transcriptional level.

### Nucleolin binding stabilizes its target transcripts

To test whether Nucleolin directly regulates target transcript stability, we performed tethering experiments. We placed increasing copies of BoxB hairpins in the 5’ UTR of a luciferase reporter and fused Nucleolin to the λN peptide (Figure 4G), which specifically binds the BoxB hairpin (De Gregorio et al., 1999; Lazinski et al., 1989). Consistent with our hypothesis, tethering Nucleolin to the 5’UTR of a luciferase reporter transcript significantly increased the intracellular abundance of that transcript in a BoxB and λN peptide dependent manner (Figure 4H), revealing that Nucleolin binding to a transcript can increase its stability. Moreover, Nucleolin tethering also significantly increased reporter gene expression even in the presence of multiple copies of BoxB hairpins, whose strong secondary structures are known to impede translational initiation in the 5’ UTR and reduce reporter expression in a copy-number dependent manner (Figure 4I) (Lytle et al., 2007). Together, these results reveal that Nucleolin binding to the 5’ UTR of target transcripts enhances transcript stability.

We hypothesized that 5’-tRF^Cys^ may facilitate recruitment of Nucleolin to its target transcripts, thereby enhancing transcript stability. To test this hypothesis, we performed RNA stability assays by quantifying transcript abundance upon 5’-tRF^Cys^ inhibition in 4T1 cells treated with the RNA Polymerase II inhibitor, 5,6-dichloro-1-beta-D-ribofuranosylbenzimidazole (DRB). Inhibition of 5’-tRF^Cys^ significantly reduced the stability of both Pafah1b1 and Mthfd1l (Figure 4J). To provide further evidence for the direct regulation of transcript stability by 5’-tRF^Cys^, we transfected *in vitro* transcribed luciferase reporter transcripts that contained either Pafah1b1 or Mthfd1l 5’ UTRs. Dual luciferase assays revealed a significant reduction in the luminescence signal upon 5’-tRF^Cys^ inhibition (Figure S4K). These data support a model in which 5’-tRF^Cys^ enhances the stability of its target transcripts by promoting Nucleolin binding to their 5’ UTRs.

### 5’-tRF^Cys^ promotes oligomerization of Nucleolin and its pro-metastatic transcripts

Given that Nucleolin directly binds both 5’-tRF^Cys^ and its target transcripts *in vivo*, we questioned how 5’-tRF^Cys^ regulates the interaction of Nucleolin with its target transcripts. A prior study reported that Nucleolin self-interacts (Chen et al., 2012). Indeed, anti-FLAG IP from cells transfected with HA-tagged Nucleolin with or without FLAG-tagged Nucleolin confirmed that Nucleolin self-interacts *in vivo* (Figure 5A). Interestingly, this self-interaction is RNA-dependent since the interaction was abolished by pre-treating lysates with RNase A before immunoprecipitation (Figure 5A). Given these findings, we hypothesized that 5’-tRF^Cys^ mediates the assembly of Nucleolin into a higher-order complex with its target transcripts.

**Figure 5.**
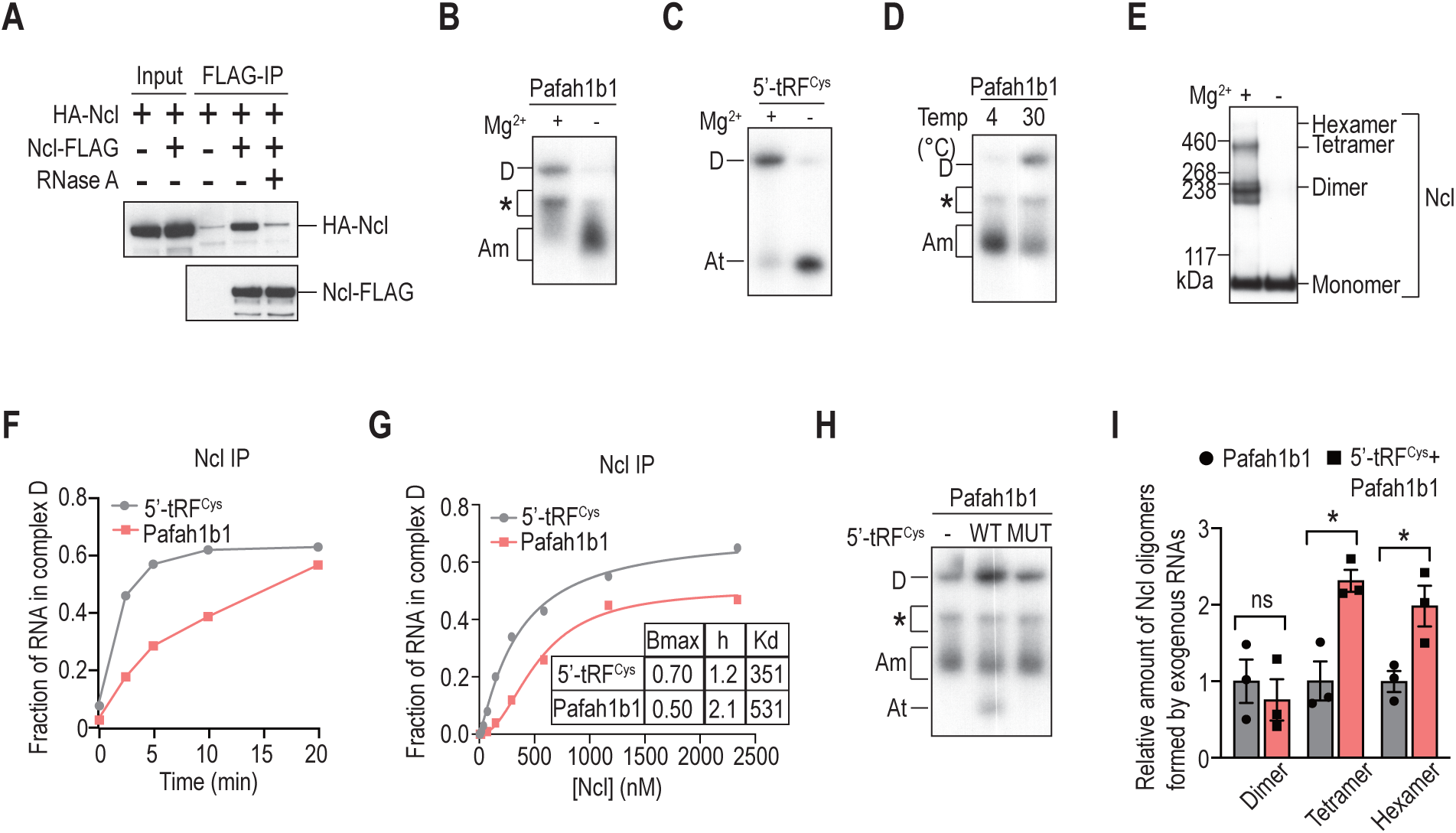
5’-tRF^Cys^ promotes complex D assembly and Nucleolin oligomerization. A. Representative images of western blots of anti-FLAG immunoprecipites (IP) from cells transfected with either HA-tagged Nucleolin alone or together with FLAG-tagged Nucleolin in the presence or absence of RNase A. B, C. Native gel analysis of complexes assembled from Pafah1b1 (B) or 5’-tRF^Cys^ (C) using Nucleolin IP in the presence or absence of Mg^2+^. Asterisk denotes an RNA-protein complex that was detected only with Nucleolin IP but not Nucleolin protein. D. Native gel analysis of complexes assembled from Pafah1b1 using Nucleolin IP at different temperatures. E. Western blot of Nucleolin using Nucleolin IP that was incubated at 30 °C with or without Mg^2+^ before crosslinking with ethylene glycol bis (succinimidyl succinate). F. Quantification of the kinetics of complex D assembly using Nucleolin IP with either Pafah1b1 or 5’-tRF^Cys^. See also Figures S5E and S5F. G. Quantification of complex D assembly using increasing amount of Nucleolin IP with either Pafah1b1 or 5’-tRF^Cys^. Bmax, specific maximum binding. h, Hill coefficient. Kd, equilibrium dissociation constant. See also Figures S5G and S5H. H. Native gel analysis of Nucleolin complexes assembled using Nucleolin IP with either Pafah1b1 alone, or together with a wild-type (WT) or a Nucleolin binding deficient 5’-tRF^Cys^ (MUT). I. Quantification of different forms of Nucleolin oligomers assembled using Nucleolin IP with Pafah1b1 alone or together with 5’-tRF^Cys^.

To test this hypothesis, we established an *in-vitro* Nucleolin complex assembly assay using 5’-radiolabeled 5’-tRF^Cys^ and/or Nucleolin’s target transcripts, with purified Nucleolin protein. We then resolved the reaction products by performing native polyacrylamide gel electrophoresis (PAGE) and detected the assembled complexes by autoradiography. In the presence of Mg^2+^, Nucleolin first formed two low-molecular-weight complexes with Pafab1b1 and 5’-tRF^Cys^, which we referred to as complex Am and At, respectively (Figures S5A and S5B), as both complexes migrated to similar positions and their assembly was independent of either Mg^2+^ or incubation at an elevated temperature. Surprisingly, as the amount of Nucleolin protein was increased, complexes Am and At gradually disappeared while a higher molecular weight complex D emerged. This suggested that complexes Am and At represent precursors of complex D (Figures S5A and S5B). Quantitative analysis of complex D assembly revealed the Hill coefficient, a measure of cooperative binding, to be 1.4 and 3.5 for 5’-tRF^Cys^ and Pafab1b1, respectively (Figure S5C), suggesting that there is cooperativity in complex D assembly from complex At for Pafah1b1 (Figure S5C).

We calculated maximum specific binding (Bmax) of complex D to be only 0.1 (Figure S5C), suggesting that formation of complex D is inefficient under the experimental condition tested. After extensive optimization, we found that complex D assembly could be substantially enhanced by using Nucleolin immunoprecipitates (IP) instead of purified Nucleolin protein, suggesting the presence of one or more unidentified factor that facilitate complex formation. We also found that complex D assembly required Mg^2+^ and incubation at an elevated temperature because its assembly was suppressed by EDTA or incubation at 4 °C (Figures 5B-5D).

We hypothesized that complex Am/At and D represent RNA-Nucleolin monomers and oligomers, respectively. To test this hypothesis, we performed chemical crosslinking and western blotting with an anti-Nucleolin antibody after the assembly assay. Consistent with this hypothesis, under experimental conditions favorable for complex D assembly (i.e., in the presence of Mg^2+^ at 30 °C), we were able to detect not only Nucleolin monomers, but also dimers, tetramers and hexamers (Figure 5E). In contrast, only monomers were detected under experimental conditions permissive for complex Am/At assembly (i.e., in the presence of EDTA) (Figure 5E). Addition of increasing amounts of micrococcal nuclease before the assembly assay gradually turned Nucleolin oligomers back into monomers (Figure S5D), supporting an essential role for RNA in promoting Nucleolin oligomerization.

Kinetic analyses revealed that 5’-tRF^Cys^ assembled complex D much faster than Pafah1b1 (Figures 5F, S5E-S5F). Quantitative analyses showed that 5’-tRF^Cys^ exhibits a lower Kd than Pafah1b1 in assembling complex D (Figure 5G, S5G-S5H). Importantly, the Hill coefficient for 5’-tRF^Cys^ and Pafab1b1 is 1.2 and 2.1 respectively (Figure 5G), suggesting Pafab1b1 exhibits much higher cooperativity than 5’-tRF^Cys^ in assembling complex D. Taken together, these results suggest that 5’-tRF^Cys^ may drive Nucleolin oligomerization with its target transcripts.

To test this hypothesis, we added wild-type or mutant 5’-tRF^Cys^ with both Nucleolin binding motifs mutated, along with its target transcripts to the assembly assay. Indeed, we found that addition of only wild-type but not mutant 5’-tRF^Cys^ could promote complex D assembly (Figures 5H and S5I). Additionally, more Nucleolin tetramers and hexamers were detected when we supplemented 5’-tRF^Cys^ along with Pafah1b1 to the assembly assay than Pafah1b1 alone (Figures 5I and S5J). Overall, these results reveal that 5’-tRF^Cys^ facilitates the oligomerization of RNA-Nucleolin monomers.

### Nucleolin oligomerization protects pro-metastatic transcripts from exonucleolytic degradation

Given that Nucleolin binds its target transcripts in either monomeric or oligomeric forms, we next assessed their functional differences. Considering that Nucleolin preferentially binds the 5’ UTR of its target transcripts, we hypothesized that oligomerization may provide protection for its target transcripts from exonucleolytic degradation. As such, we tested the sensitivity of Nucleolin’s target transcripts to a 5’->3’ exonuclease under experimental conditions permissive for Nucleolin oligomerization. Remarkably, we found that target transcripts were protected by Nucleolin from degradation by the 5’->3’ exonuclease when the complex assembly assay was performed under experimental conditions favorable for its oligomerization (i.e., at 30 °C) in comparison to conditions favoring only monomeric Nucleolin (i.e., at 4 °C) (Figure S5K). Thus, in comparison to monomeric Nucleolin, oligomeric Nucleolin provides better protection for its target transcripts from exonucleolytic degradation.

### Pafah1b1 and Mthfd1l function downstream of 5’-tRF^Cys^ to promote breast cancer metastasis

Having shown that expression of both Pafah1b1 and Mthfd1l is modulated by 5’-tRF^Cys^, we questioned whether they were the downstream effectors of the pro-metastatic phenotype of 5’-tRF^Cys^. We thus performed epistasis experiments. Overexpression of Pafah1b1 or Mthfd1l rescued survival defects mediated by 5’-tRF^Cys^ inhibition *in vivo*, as quantified by cleaved Caspase 3/7 activity (Figures S6A and S6B). Additionally, overexpression of either Pafah1b1 or Mthfd1l rescued metastatic lung colonization defects upon 5’-tRF^Cys^ inhibition (Figures 6A and 6B), consistent with these genes functioning downstream of 5’-tRF^Cys^ in promoting breast cancer metastasis. Together, these results reveal that Pafah1b1 and Mthfd1l enhance breast cancer cell survival and are downstream effectors of 5’-tRF^Cys^-mediated metastatic colonization.

**Figure 6.**
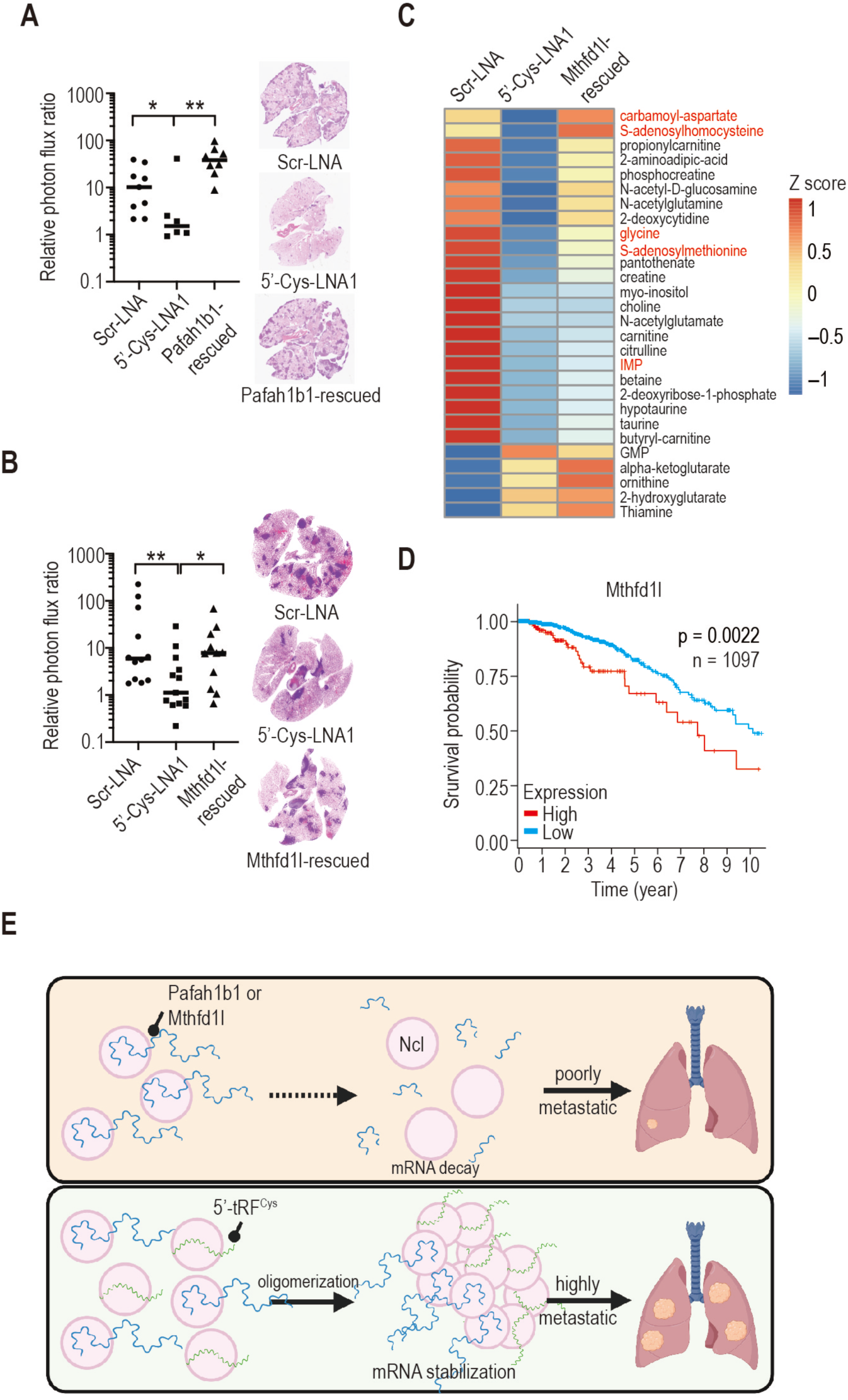
Pafah1b1 and Mthfd1l function downstream of 5’-tRF^Cys^ to promote breast cancer metastasis. A, B. Bioluminescence quantification of metastatic lung colonization in mice injected with 4T1 cells transfected with a control LNA (Scr-LNA), a 5’-tRF^Cys^ targeting LNA (5’-Cys-LNA1) either alone or together with overexpression of Pafah1b1 (Pafah1b1-rescued) (A) or Mthfdl1 (Mthfd1l-rescued) (B). Representative H&E staining of lung sections for each cohort are shown (N=6-13). C. Heatmap showing abundance z-scores for metabolites that were significantly changed upon 5’-tRF^Cys^ inhibition in 4T1 cells. D. Kaplan-Meier curves depicting survival probability of breast cancer patients in the TCGA cohort (n=1097) stratified by the expression of Mthfd1l. Statistical significance was determined by Mantel–Cox log-rank test. E. A schematic model depicting the mechanism underlying 5’-tRF^Cys^ promotion of breast cancer metastasis.

Pafah1b1, together with two catalytic enzymes Pafah1b2 and Pafah1b3, form the type I PAF-AH complex (Arai et al., 2002). Pafah1b1 is proposed to control the level of platelet activating factor (PAF) by regulating the enzymatic activity of Pafah1b2 and Pafah1b3 (Arai et al., 2002). Consistent with our finding, prior studies reported that Pafah1b2 and Pafah1b3 promote cancer progression and metastasis (Ma et al., 2018; Mulvihill et al., 2014). On the other hand, Mthfd1l is a key metabolic enzyme in the folate cycle (Figure S6C) and has been recently reported to promote progression of several cancer types (Agarwal et al., 2019; Eich et al., 2019; Lee et al., 2017). Moreover, Mthfd1l was previously found to promote breast cancer cell survival and its expression was observed to become elevated in highly metastatic breast cancer cells (Li et al., 2020; Selcuklu et al., 2012). To search for Mthfd1l downstream metabolite effectors that may promote metastasis, we performed untargeted metabolite profiling. Consistent with our finding that Mthfd1l is the downstream effector of 5’-tRF^Cys^ and the well-established role of Mthfd1l in one-carbon metabolism (Ducker et al., 2016; Ducker and Rabinowitz, 2017; Jain et al., 2012; Minton et al., 2018; Zheng et al., 2018), we identified several metabolites in the folate cycle (glycine) and interconnected metabolomic pathways (e.g. inosine monophosphate (IMP), carbamoyl-aspartate (CA), S-adenosylhomocysteine (SAH) and S-adenosylmethionine (SAM)), that became significantly increased in both control and Mthfd1l-rescued cells relative to the 5’-tRF^Cys^ inhibited cells (Figures 6C and S6C). Moreover, we found that supplementation with both formate and glycine, two key metabolites of the folate cycle, rescued the cell survival defect in 5’-tRF^Cys^-suppressed cells (Figure S6D). These results collectively reveal that 5’-tRF^Cys^ regulates one-carbon metabolism through Mthfd1l, thereby contributing to metastasis.

### Elevated expression of 5’-tRF^Cys^ targets correlate with poor clinical outcome in breast cancer patients

We next sought to determine if the expression levels of 5’-tRF^Cys^ target transcripts Pafah1b1 and Mthfd1l associate with clinical outcomes of human breast cancer patients. Breast cancer patients whose tumors expressed higher levels of Pafah1b1 and Mthfd1l exhibited significantly reduced survival relative to those whose tumors expressed lower levels of these genes (Figures 6D and S6E; p<0.0001 and p=0.0022, respectively; n=1097). Together, these findings provide human pathological association support for a role for this tRF/Pafah1b1/Mthfd11 axis in post-transcriptional control of metabolic response in driving breast cancer progression.

## DISCUSSION

*In vivo* and *in vitro* experiments reveal that 5’-tRF^Cys^ promotes metastatic lung colonization of breast cancer cells by enhancing cancer cell survival. We find that two metabolic enzymes—Pafah1b1 and Mthfd1l— act downstream of 5’-tRF^Cys^ to enhance cancer cell survival and metastasis. Our findings are consistent with the emerging recognition of the impact of post-transcriptional regulation on metabolic control under pathological conditions (Loo et al., 2015; Sullivan et al., 2018; Zhu et al., 2011), and underscore the critical role that dysregulated metabolism plays in tumor biology (Cantor and Sabatini, 2012; DeBerardinis and Chandel, 2016; Pavlova and Thompson, 2016).

While 5’-tRF^Cys^ was reported to bind YBX-1 in a study using rabbit reticulocyte lysates (Ivanov et al., 2011), we find that Nucleolin is the primary binding partner of 5’-tRF^Cys^ *in vitro* and *in vivo* in breast cancer cells. HITS-CLIP analysis and EMSA assays revealed that Nucleolin binds 5’-tRF^Cys^ by engaging its two G-rich motifs. A prior study using Selection of Ligands by Exponential Enrichment (SELEX) proposed that Nucleolin binds RNAs through an evolutionary conserved motif (Ginisty et al., 2000). Interestingly, upon close inspection of the selected sequences, we identified the enrichment of a highly similar G-rich motif from this past study. We speculate that the same G-rich motif identified herein may contribute to Nucleolin binding to the SELEX-enriched sequences in that prior study.

Although Nucleolin is best known for its role in rRNA biogenesis and ribosome assembly, we found that Nucleolin binds a previously unappreciated range of RNAs. Interestingly, the majority of binding sites in mRNAs reside within 5’ UTRs, suggesting a direct role for Nucleolin in post-transcriptional regulation. In addition to the well-established role for RNA binding proteins in regulating the stability of transcripts via 3’ UTR-binding, recent studies have uncovered a similar role for RNA binding proteins at 5’ UTR sites (Arribere and Gilbert, 2013; Leppek et al., 2018). Indeed, tethering of Nucleolin to the 5’ UTR increased the abundance of mRNA transcripts *in vivo*. Additionally, complex D, which represents an RNA-Nucleolin oligomer, better protected RNAs from exonucleolytic degradation relative to an RNA-Nucleolin monomer *in vitro*. Together, these results are consistent with a model whereby Nucleolin oligomerization plays a critical role in stabilizing its bound target transcripts. While we found that Nucleolin binding at the 5’ UTR primarily enhances transcript stability under our experimental conditions, we cannot exclude the possibility that Nucleolin binding may play additional roles in post-transcriptional regulation of mRNAs under different experimental conditions, as previously reported (Abdelmohsen et al., 2011; Izumi et al., 2001; Takagi et al., 2005).

Our data support a model wherein 5’-tRF^Cys^ aids in the formation and stabilization of higher order mRNA-Nucleolin complexes. 5’-tRF^Cys^ exhibits a higher affinity for Nucleolin and assembles complex D at a faster rate than its target transcripts, while target transcripts display higher binding cooperativity in Nucleolin oligomerization than 5’-tRF^Cys^, suggesting 5’-tRF^Cys^ may act as a nucleator for oligomerization from monomeric mRNA-Nucleolin complexes. Given that both complex D assembly and Nucleolin oligomerization require Mg^2+^ and incubation at an elevated temperature, and that Nucleolin IP is more efficient at oligomerization than purified Nucleolin protein, it is tempting to speculate that one or more enzymes that interact with Nucleolin may catalyze complex D assembly and Nucleolin oligomerization. Future studies are required to identify the enzyme(s) that are responsible for this transformation.

Although we observed that the oligomeric RNA-Nucleolin complex better protects its target transcripts from 5’->3’ exonucleolytic decay than the monomeric RNA-Nucleolin complex, it remains possible that the oligomeric RNA-Nucleolin complex may additionally stabilize its target transcripts by suppressing decapping and/or 3’->5’ exonucleolytic degradation. It is intriguing to note that oligomeric Nucleolin was detected in even numbers of units (dimers, tetramers or hexamers) but not odd numbers of units. Therefore, it is possible that Nucleolin dimers, rather than monomers, may be the building blocks for assembling the oligomeric complex. While we were able to detect up to hexameric forms of Nucleolin, we cannot exclude the possibility that oligomeric complexes of higher orders may exist. Defining and characterizing the molecular components and functional interactions of this oligomeric Nucleolin complex will require future biochemical and structural studies.

## METHODS

### Cell Culture

67NR, 4TO7 cells were generously provided by W. P. Schiemann (Case Comprehensive Cancer Center). 4T1 cells were obtained from ATCC. MDA-MB-231-LM2 (Minn et al., 2005), 67NR, 4TO7 and 4T1 cells were maintained in DMEM supplemented with 10% FBS, sodium pyruvate, L-glutamine, Pen/Strep and Amphotericin. EO771 and its metastatic lung derivative EO771-LM3 were cultured in RPMI supplemented with 10% FBS, sodium pyruvate, L-glutamine, Pen/Strep, Amphotericin and 10 mM HEPES. To inhibit 5’-tRF^Cys^, 50 nM of scrambled control or 5’-tRF^Cys^ antisense locked nucleic acid (LNA) oligos were transfected into cells with Lipofectamine 2000 (Thermo Fisher Scientific). Experiments were performed 24-48 hours post transfection. All cell lines were routinely tested for the presence of mycoplasma and were confirmed to be free of mycoplasma contamination.

### Patient-Derived Xenograft Organoids

Breast cancer patient-derived xenograft organoids (PDXO) were obtained from A. L. Welm (DeRose et al., 2011; Sikora et al., 2014). Organoids were embedded in reduced growth factor matrigel (BD Biosciences) and the growth media was made from advanced DMEM/F12 (Thermo Fisher Scientific) supplemented with 5% FBS (Thermo Fisher Scientific), 10 mM Hepes, GlutaMAX (Thermo Fisher Scientific), 1 ug/ml of Hydrocortisone, 0.05 mg/ml of Gentamicin and 10 ug/ml hEGF. This medium was kept at 4°C for up to 2 weeks and prior to addition to PDXO cultures supplemented with 10μM Rock Inhibitor Y-27632 (Selleck Chemicals), 100 ng/ml FGF2 (R&D Systems), 1 mM N-Acetyl-L-cysteine (Sigma Aldrich) and 10 mM Heregulin Beta-1 (PeproTech).

PDXO cells were transduced with pHIV-Luc-ZsGreen (a gift from B. Welm; Addgene# 39196), pLKO.1-Scr-TD or pLKO.1-Cys-TD as previously described (Tavora et al., 2020). Lentivirus was concentrated using Lenti-X Concentrator (Takara) and re-suspended in PDXO culture media supplemented with 10 μg/ml of polybrene (Sigma Aldrich). Organoids were dissociated and resuspended in the PDXO culture media previously used to re-suspend the virus and transferred to a 24 well ultra-low attachment plate (Corning) and centrifuged at 600 g for 1 hour at room temperature and then incubated at 37°C for 6 hours. Organoids were expanded in matrigel for another two weeks and then zsGreen positive organoids were sorted using a BD FACSAria II cell sorter and expanded again in matrigel. Alternatively, organoids were selected with 1 ug/ml Puromycin until all non-transduced organoids were dead.

### Mouse Experiments

4T1, EO771-LM3, MDA-MB-231-LM2 and PDXO cells that have been labeled with a luciferase and GFP reporter or a luciferase and ZsGreen reporter were injected into 6-10 week-old female Balb/c, C57Bl/6 and immunocompromised NOD-scid-gamma (NSG) mice respectively via tail vein. Tumor growth was monitored weekly by luminescence imaging. Lung nodules were detected by H&E staining.

### Molecular Cloning

FLAG-tagged mouse Nucleolin cDNA was cloned into pcDNA3. Mouse Mthfd1l or Pafah1b1 cDNAs were cloned into pLJC6-EF1core-Blast vector. The 5’ UTR of mouse 5’-tRF^Cys^ targets or non-target Gapdh were cloned into psiCHECK2 (Promega). Synthetic 5’-tRF-Cys antisense or a scrambled control sequence embedded in the tough decoy backbone (Haraguchi et al., 2009; Xie et al., 2012) were cloned into the pLKO.1-Puro vector (a gift from R. Weinberg; Addgene #8453) to generate pLKO.1-Cys-TD and pLKO.1-Scr-TD, respectively.

### RNA Isolation

Unless stated otherwise, RNAs were isolated using TRIzol and RNeasy MINI kit (Qiagen) with the following modifications to isolate both large (>200 nt) and small (<200 nt) RNAs. In brief, chloroform-extracted lysates were supplemented with the addition of 1.5 volumes of ethanol before lysates were passed through the column and the column was washed with only buffer RPE twice.

### Lentiviral transduction

Lentiviral viruses were produced as described previously (Liu et al., 2014). In brief, 293LTV cells were co-transfected with pLKO.1-Puro-Scr-TD, pLKO.1-Puro-Cys-TD, pLJC6-EF1core-Blast-mPafah1b1, or pLJC6-EF1core-Blast-mMthfd1l, plus pSPAX2 and pMD2.G (gifts from D. Trono; Addgene #12260 and #12259) using Lipofectamine 2000. After 48 hours, lentiviruses were concentrated using Lenti-X concentrator (Takara). 4T1 or MDA-MB-231 cells were transduced with concentrated viruses in the presence of 8 ug/ml of polybrene (Sigma) overnight or spin transduced at 500g at 32 °C for 90 min followed by incubation at 37 °C for 6 hours. Transduced cells were selected with 2-3 ug/ml of Puromycin or 3 ug/ml of Blasticidin 2 days post transduction until all non-transduced cells were dead.

### Cell Viability and Apoptosis Assays

Cells were seeded in a 24-well plate and transfected with a control or 5’-tRF^Cys^ antisense LNA as described above, cell viability and Caspase activity were detected with CellTiter-Glo 2.0 and CaspaseGlo-3/7 assay (Promega) respectively 2 days after transfection. Caspase 3/7 activity was determined by dividing the CaspaseGlo-3/7 signal by the CellTiter-Glo 2.0 signal. Alternatively, cells were seeded in a 6-well plate and transfected with a control or 5’-tRF^Cys^ antisense LNA as described above, fraction of apoptotic cells were determined using PE Annexin V Apoptosis Detection Kit I (BD Bioscience) with an Attune Nxt Flow Cytometer (Thermo Fisher Scientific).

### Immunostaining

Immunostaining of mouse lung sections was performed as described previously (Tavora et al., 2020) with the following antibodies: anti-Ki-67 (Abcam), anti-Endomucin (Santa Cruz Biotechnology), anti-cleaved Caspase3 (Asp175) (Cell Signaling Technology), anti-rabbit Alexa Fluor 488 and anti-rat Alexa Fluor 555 (Thermo Fisher Scientific). Images were acquired using a Zeiss LSM 880 confocal microscope. H&E staining of mouse lung sections were performed by Histoserv Inc.

### CLIP RT-qPCR

4T1 cells were lysed with lysis buffer (1×PBS (no Mg^2+^ or Ca^2+^), 0.1% SDS, 0.5% deoxycholate and 0.5% IGEPAL CA-630) plus cOmplete, EDTA-free protease inhibitor and 0.5 U/ul RNasin Plus RNase inhibitor. 40 ul of RQ1 DNase I (Promega) was added, and lysate was shaken at 37 °C at 1,000 rpm for 10 min. The lysate was spun at 16,000 g at 4 °C for 10 min. The supernatant was transferred to a fresh tube. Dynabeads M-280 Sheep Anti-Rabbit IgG (Thermo Fisher Scientific) coupled to either a rabbit IgG control or an anti-Nucleolin antibody (Cell Signaling Technology) was added and rotated in the cold room for 1 hour. The supernatant was discarded, and beads were washed twice with lysis buffer, twice with high salt wash buffer (5×PBS (no Mg^2+^ or Ca^2+^), 0.1% SDS, 0.5% deoxycholate, 0.5% IGEPAL CA-630) and twice with PNK buffer (50 mM Tris-HCl, pH 7.5, 10 mM MgCl_2_, 0.5% IGEPAL CA-630). The beads were drained of all liquids before 2 mg/ml of Proteinase K (New England Biolabs) diluted in PK buffer (50 mM Tris-HCl pH 7.5, 10 mM MgCl_2_, 0.5% IGEPAL CA-630) plus 0.2% SDS was added and shaken at 1,000 rpm at 50 °C for 30 min. RNAs were purified sequentially with acid phenol-chloroform (Thermo Fisher Scientific) and chloroform before precipitated overnight with ethanol. Precipitated RNAs were quantified using Taqman miRNA assays with custom primers for detection of 5’-tRF^Cys^ (Thermo Fisher Scientific).

### Small RNA Sequencing and Analysis

Libraries were constructed from total RNAs isolated from 67NR, 4TO7 or 4T1 cells using RealSeq-AC (SomaGenics) according to the manufacturer’s instruction. All high-throughput sequencing libraries were sequenced for 75 cycles using NextSeq 500 or 150 cycles using HiSeq 2500 at the Rockefeller University Genomics Resource Center. Adapters were removed from the 3’ end of reads using cutadapt (Martin, 2011). To identify differentially expressed tRNA fragments, trimmed reads were aligned using bwa (Li and Durbin, 2009) to the mouse mature tRNA space, which consists of all mouse tRNA sequences downloaded from GtRNAdB (Chan and Lowe, 2009, 2016) with introns and CCA-tail removed and identical sequences collapsed. Mapped reads were counted using featureCounts (Liao et al., 2014) and differentially expressed tRNA fragments were determined using DESeq2 (Love et al., 2014).

### RNA Sequencing and Analysis

Total RNA was depleted of ribosomal RNAs using NEBNext rRNA Depletion Kit (Human/Mouse/Rat) (New England Biolabs) before they were used for library construction using NEBNext Ultra II Directional RNA Library Prep Kit (New England Biolabs). Adaptors were removed from the 3’ end of reads using cutadapt (Martin, 2011). Trimmed reads were aligned to the mouse genome (mm10) using STAR (Dobin et al., 2013). Uniquely mapped reads were counted using featureCounts (Liao et al., 2014) and differentially expressed genes were determined using DESeq2 (Love et al., 2014).

### HITS-CLIP and Analysis

Regular HITS-CLIP libraries were constructed as described previously (Moore et al., 2014). To identify small RNAs directly bound by Nucleolin, HITS-CLIP was performed without use of any RNase. Analysis was done using CLIP Tool Kit (Shah et al., 2017). Piranha was used to identify Nucleolin-bound peaks with a background threshold of 0.95 (Uren et al., 2012). The identified peaks from all samples were merged before reads that aligned to the peaks were counted using featureCounts (Liao et al., 2014).

### Proteomic Analysis

4T1 cells transfected with a scrambled control or 5’-tRF^Cys^ antisense LNA were lysed in lysis buffer (20 mM Tris-HCl, pH 7.5, 100 mM KCl, 1 mM EDTA, 0.5% NP-40) supplemented with cOmplete EDTA-free protease inhibitor and 1 mM DTT. Lysates were sonicated 10 seconds for three times and centrifuged at 16,000 g at 4 °C for 10 minutes. The supernatant was sent to identify differentially expressed proteins by reversed phase nano-LC-MS/MS (Ultimate 3000 coupled to a Q-Exactive HF, Thermo Scientific) at the Rockefeller University Proteomics Resource Center.

### Ribosome Profiling and Analysis

Ribosome profiling (Ribo-Seq) libraries were constructed as described previously (McGlincy and Ingolia, 2017). Adaptors were removed from the 3’ end of reads using cutadapt (Martin, 2011). Trimmed reads were aligned to the mouse genome (mm10) using STAR (Dobin et al., 2013). Both uniquely- and primary multi-mapped reads that aligned to the annotated coding sequence of protein coding genes were counted using featureCounts (Liao et al., 2014) and differentially translated genes were determined using DESeq2 (Love et al., 2014).

### Radiolabling of Nucleolin Target Transcripts and 5’-tRF^Cys^

DNA templates for Nucleolin-bound regions were PCR amplified and in-vitro transcribed with MEGAshortscript T7 transcription kit (Thermo Fisher Scientific). In-vitro transcribed products and chemically synthesized 5’-tRF^Cys^ (Integrated DNA Technologies) were purified on a 10% TBE-Urea PAGE gel (Thermo Fisher Scientific) before 5’ radiolabeled using T4 PNK (New England Biolabs). Labeling reactions were purified with acid-phenol chloroform and chloroform before precipitated overnight with ethanol.

### Electrophoretic Mobility Shift Assay

Purified Nucleolin protein was incubated with 5’ radiolabeled 5’-tRF^Cys^ and/or in-vitro transcribed 5’-tRF^Cys^ targets in binding buffer (20 mM HEPES, pH 7, 50 mM KCl, 1 mM DTT, 1 mM EDTA) at 4 °C for 10 min before 3% of Ficoll-400 (Sigma) was added and assembled Nucleolin complexes were resolved on a 10% TBE PAGE gel at 4 °C.

### Dual Luciferase Assay

pcDNA3-FLUC containing zero, two or five BoxB sites were constructed from phRL-TK-5BoxB-sp36 (a gift from W. Filipowicz; Addgene #115365) (Pillai et al., 2004) and pcDNA3-RLUC-POLIRES-FLUC (a gift from N. Sonenberg; Addgene #45642) (Poulin et al., 1998) using Gibson assembly. HA-mNcl and λNHA-mNcl were constructed by subcloning mNcl from pcDNA3-mNcl-FLAG into pCIneo-deltaHA and pCIneo-deltaNHA (gifts from W. Filipowicz; Addgene #115360, #115359) (Pillai et al., 2004). Firefly luciferase reporters and λNHA-LacZ (a gift from W. Filipowicz; Addgene #115363) (Pillai et al., 2004), HA-mNcl or λNHA-mNcl were co-transfected with pNL1.1.CMV (Promega) into unlabeled 4T1 cells using Lipofectamine 2000 (Life Technologies). Dual luciferase assay was performed 2 days post transfection using the Nano-Glo Dual-Luciferase Reporter Assay (Promega) in a Synergy Neo2 microplate reader (Biotek). Alternatively, 5’ UTR of Renilla luciferase gene was fused with Nucleolin-bound regions from the target transcript or GAPDH. Renilla luciferase gene and Firefly luciferase gene were in-vitro transcribed using HiScribe T7 RNA Synthesis Kit (NEB Biolabs). After purified with Monarch RNA Cleanup Kit (NEB Biolabs), RNAs were polyadenylated with E. coli poly(A) polymerase (NEB Biolabs), and capped and 2’-O-methylated with Vaccinia Capping System (NEB Biolabs). LNA oligos were transfected into unlabeled 4T1 cells using Lipofectamine 2000 (Thermo Fisher Scientific) as described above. The next day, Firefly and Renilla luciferase reporter transcripts were co-transfected into unlabeled 4T1 cells using Lipofectamine RNAiMax (Thermo Fisher Scientific) for 6-8 hours before dual luciferase assay was performed using the Dual-Glo Luciferase Assay (Promega) in a Synergy Neo2 microplate reader (Biotek).

### Metabolite Profiling

Cells cultured in 6-well plates were transfected with 50 nM of scrambled control or 5’-tRF-Cys antisense LNA oligos using Lipofectamine 2000 as described above. After 24 hours, cells were washed with 0.9% NaCl twice and lysed with pre-chilled extraction buffer that was composed of 480 ul of MeOH mixed with 120 ul of H2O and 1 uM of pre-mixed Heavy Amino Acid mix (Cambridge Isotope Laboratories). Lysates were transferred to an Eppendorf tube and vortexed at 4 °C for 10 min before being centrifuged at 16,000 g at 4 °C for 10 min. Supernatant was transferred to a fresh Eppendorf tube and dried using a nitrogen air evaporator. The dried pellet was analyzed by LC-MS/MS at the Rockefeller University Proteomics Resource Center. The pellets were resuspended in 60 μl of 50% acetonitrile, vortexed for 10 seconds, centrifuged for 15 minutes at 20,000 g at 4°C and 5 μl of the supernatant was injected onto the LC-MS in a randomized sequence. Chromatographic separation and mass spectrometry analysis was acquired as previously described (Soula et al., 2020). Relative quantification of polar metabolites was performed using Skyline (Pino et al., 2020) with the maximum mass and retention time tolerance set to 2 ppm and 12 s, respectively, referencing an in-house library of polar metabolite standards.

## ACKNOWLEDGEMENTS

We thank K. Sawicka and R. Darnell for their help with polysome profiling experiments. We thank C. Zhao, C. Lai, and N. Nnatubeugo at the Genomics Resource Center, S. Mazel and the Flow Cytometry Center, the Bio-Imaging Center, the High-Throughput Screening Resource Center and the Comparative Biology Center for their technical support. We thank K. Birsoy and S. Kurdistani for their critical comments on our manuscript. We are also grateful for members of the Tavazoie lab for their helpful discussion. X.L. was supported by a Bristol Meyers Squibb postdoctoral fellowship. The research of S.F.T. was supported in part by a Faculty Scholar grant from the HHMI, by the DOD Collaborative Scholars and Innovators Award (W81XWH-12-1-0301), Pershing Square Sohn Cancer Research Alliance award, Breast Cancer Research Foundation award, Emerald Foundation, NIH grant 5R01CA215491-05, and the Black Family Metastasis Center.

## AUTHOR CONTRIBUTIONS

X.L. and S.F.T conceived and designed the project and supervised all experiments. X.L. designed and performed experiments and conduced bioinformatic and statistical analyses. X.L. and H.A. performed metabolite profiling. X.L. and H.M. performed proteomic analysis. B.T. provided technical support for PDXO experiments. X.L. and S.F.T. wrote the manuscript with input from all authors.

## Supplementary Figure Legends

**Figure 1.**
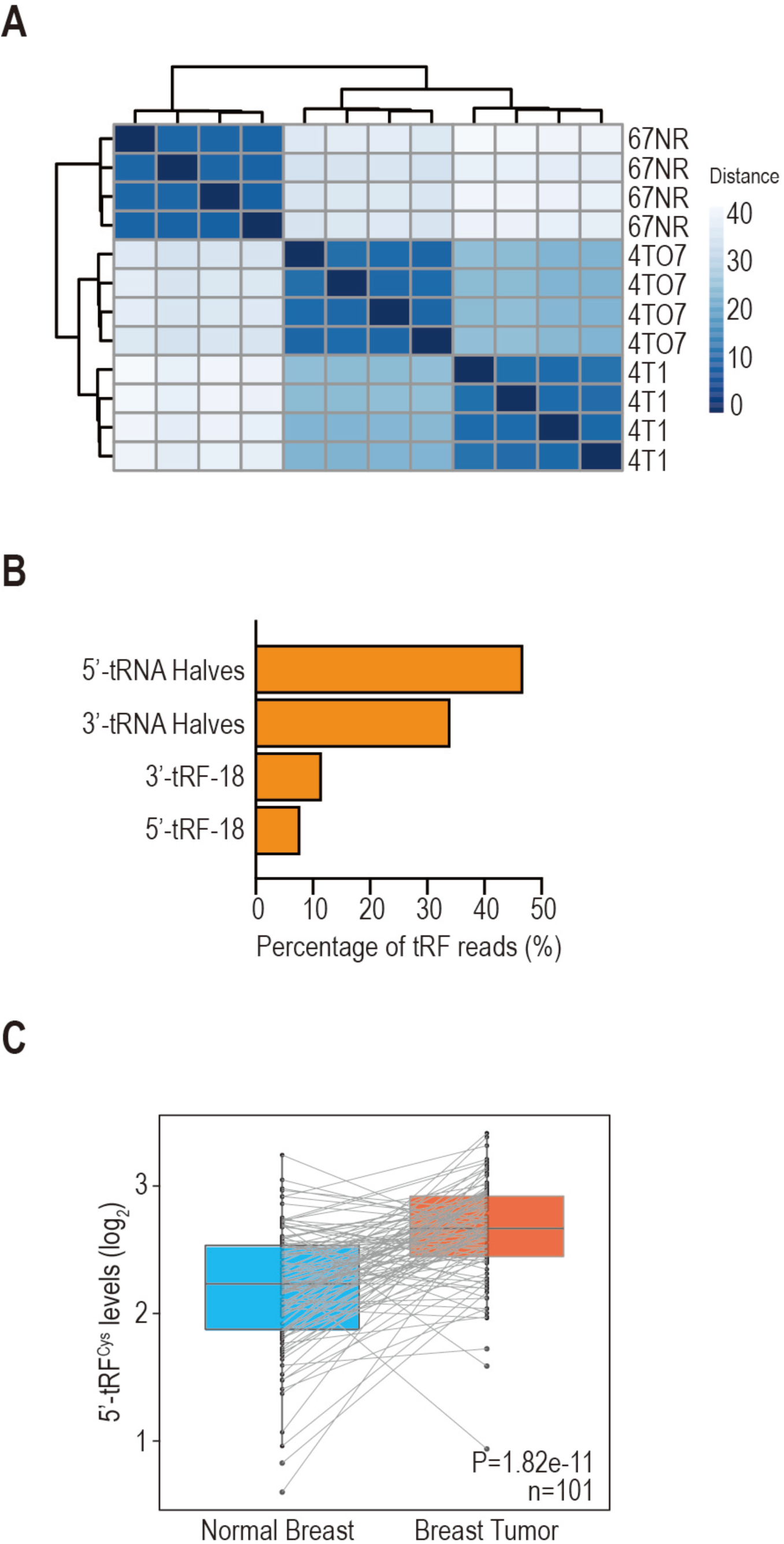
5’-tRF^Cys^ is upregulated during breast cancer progression and metastasis. A. Sample distance matrix as determined by expression of small RNAs comprised of tRNA fragments and miRNAs. B. Fraction of different types of tRNA Fragments (tRFs) in 4T1 cells. C. Expression of 5’-tRF^Cys^ in breast tumors and matched normal breast tissues in the TCGA cohort (n=101). Statistical significance was determined by paired t test.

**Figure 2.**
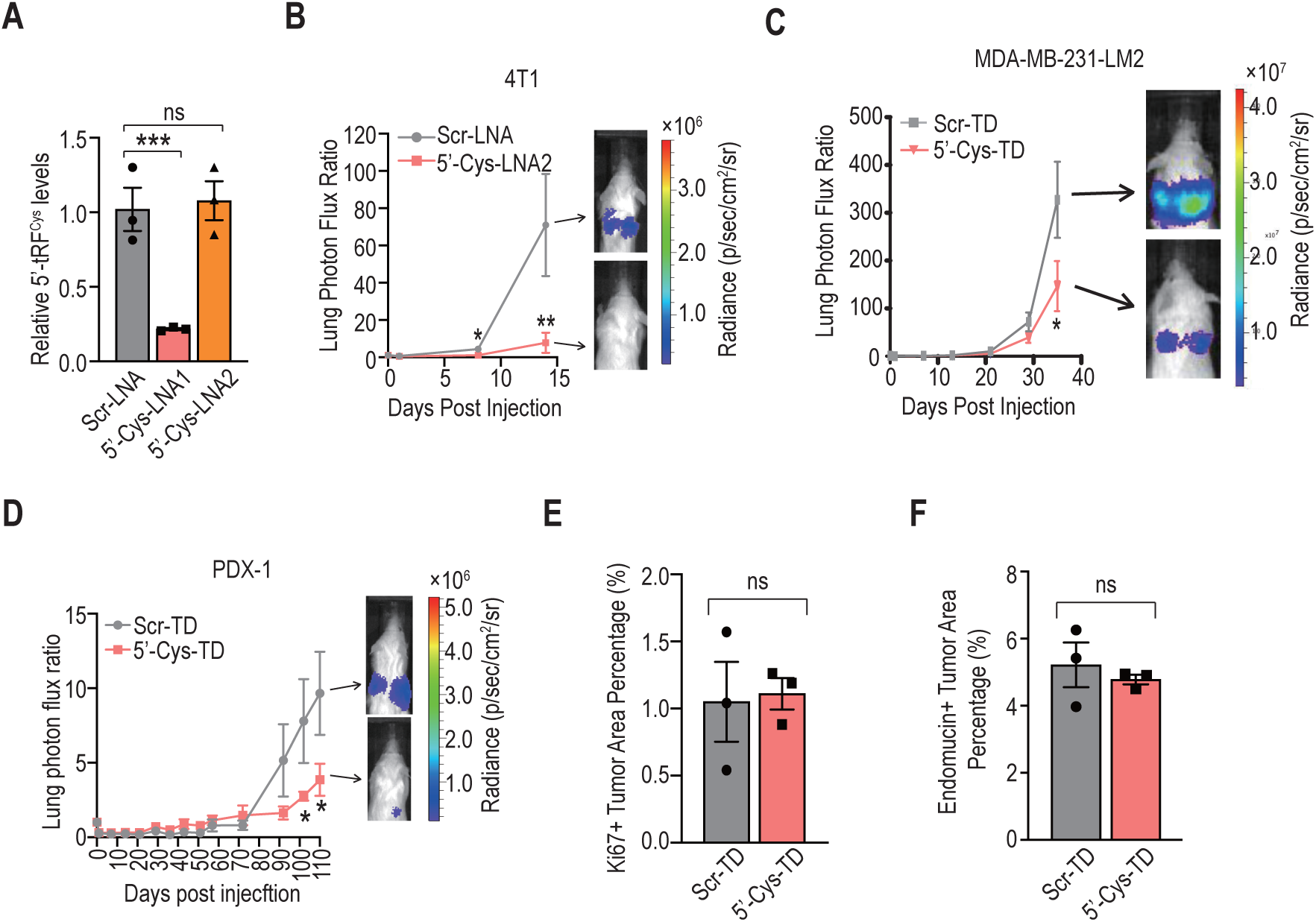
5’-tRF^Cys^ promotes breast cancer metastasis and enhances breast cancer cell survival. A. Quantification of 5’-tRF^Cys^ levels by stem-loop RT-qPCR in 4T1 cells transduced with a scramble control (Scr-LNA) or two distinct 5’-tRF^Cys^ antisense LNA oligos. All data hereafter are represented as mean ± s.e.m. and data points represent biological replicates. Representative results from at least two independent experiments are shown, except for mouse experiments, which were performed only once with cohorts of mice. B. Bioluminescence imaging plot of metastatic lung colonization in Balb/cJ mice by 4T1 cells transfected with a scramble control (Scr-LNA) or a 5’-tRF^Cys^ antisense LNA (5’-Cys-LNA2). Representative bioluminescence images for each cohort are shown (N=5-7). C, D. Bioluminescence imaging plot of metastatic lung colonization in Nod scid gamma (NSG) mice by MDA-MB-231-LM2 (C) and human breast cancer patient-derived xenograft organoid (PDXO) (D) cells transduced with a scramble control (Scr-TD) or a 5’-tRF^Cys^ antisense tough decoy molecule (5’-Cys-TD). Representative bioluminescence images for each cohort are shown (N=5-6). E, F. Percentage of area in metastatic tumor sections that are Ki-67 (E) or Endomucin (F) positive. Mice were injected with 4T1 cells transduced with a 5’-tRF^Cys^ antisense (5’-Cys-TD) or a scrambled control tough decoy (Scr-TD) (N=3). Each data point represents the average of at least 10 different image measurements from one mouse lung section. All data are represented as mean ± s.e.m. Data points represent biological replicates. Statistical significance in mouse (B-D) and molecular (A, E-F) experiments was determined by one-tail Mann-Whitney tests and t-tests with Welch’s correction respectively. ns, not significant. *, p<0.05; **, p<0.01; ***, p<0.001.

**Figure 3.**
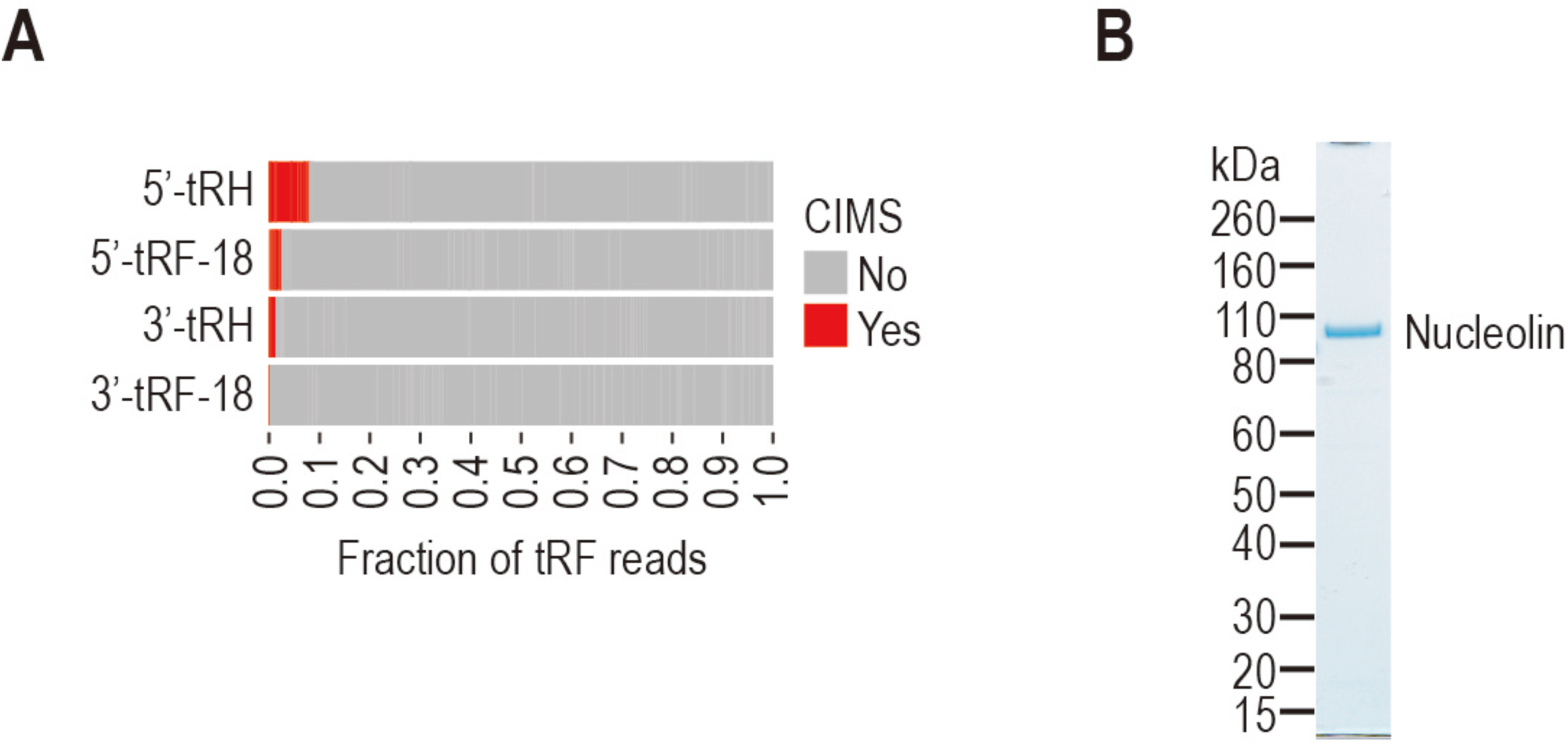
Nucleolin is a direct binding partner of 5’-tRF^Cys^. A. Fraction of different types of Nucleolin-bound tRFs that contain CIMS. B. Coomassie staining of purified Nucleolin protein.

**Figure 4.**
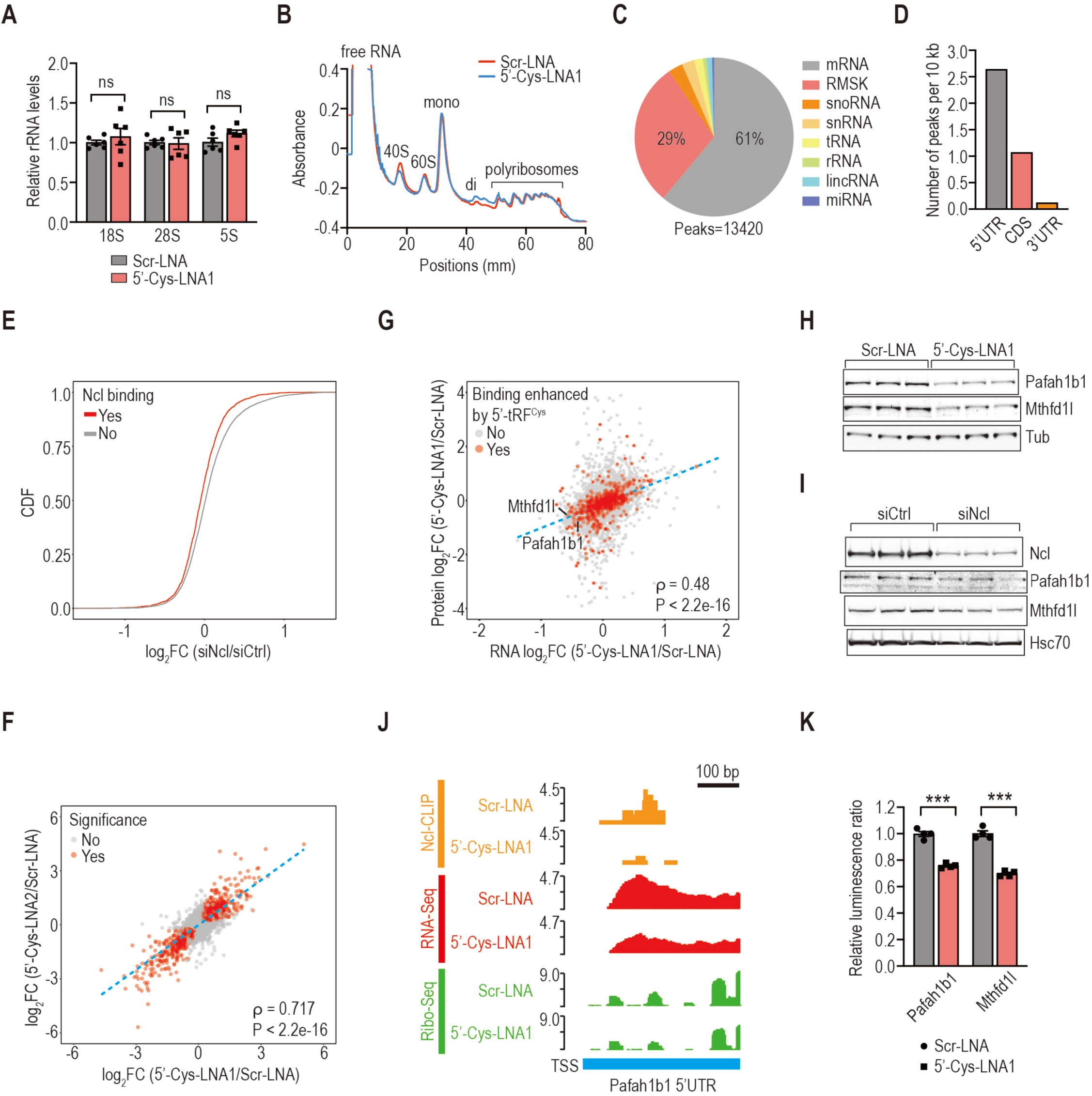
5’-tRF^Cys^ promotes Nucleolin binding to its target transcripts to enhance their stability. A. Quantification of rRNA levels upon inhibition of 5’-tRF^Cys^ by RT-qPCR. B. Representative polysome profiles showing global translation status in 4T1 cells upon inhibition of 5’-tRF^Cys^. Mono, monosomes. Di, disomes. C. Percentage of Nucleolin peaks in different types of RNAs. RMSK, repeat masked RNAs. D. The number of Nucleolin-bound CLIP peaks in 5’, 3’ untranslated region (UTR) or coding sequencing (CDS) per 10 kb in the mouse genome. E. Cumulative distribution function (CDF) plots of log_2_FC in transcript abundance for all transcripts stratified by whether they were bound by Nucleolin (red) or not (grey). Statistical significance was determined by Kolmogorov–Smirnov (KS) test (P = 4.8e-13). F. Scatter plot comparing log_2_FC in transcript abundance upon inhibition of 5’-tRF^Cys^ with two distinct 5’-tRF^Cys^ antisense LNAs. Statistically significantly changed genes are marked in red. The blue dashed line represents the linear regression line for all data points. ρ, Spearman’s correlation coefficient. G. Scatter plot comparing log_2_FC in protein abundance and log_2_FC in transcript abundance between 5’-tRF^Cys^ suppressed and control cells for all transcripts stratified by whether their Nucleolin binding is enhanced by 5’-tRF^Cys^ (red) or not (grey). The blue dashed line represents the linear regression line for all data points. ρ, Spearman’s correlation coefficient. H, I. Representative western blot images of 5’-tRF^Cys^ targets upon suppression of 5’-tRF^Cys^ (H) or depletion of Nucleolin (I). J. Genome browser view of the aligned Nucleolin (Ncl)-CLIP tags (orange), RNA-Seq reads (red) and Ribo-Seq reads (green) within the 5’ UTR of Pafah1b1. The Y axis represents reads per million (RPM). TSS, transcription start site. K. Quantification by dual luciferase assays of the luminescence signals of reporters containing 5’ UTRs from 5’-tRF^Cys^ targets relative to that from the control GAPDH. Statistical significance in A and K was determined by one-tail t-tests with Welch’s correction. ns, not significant. ***, p<0.001.

**Figure 5.**
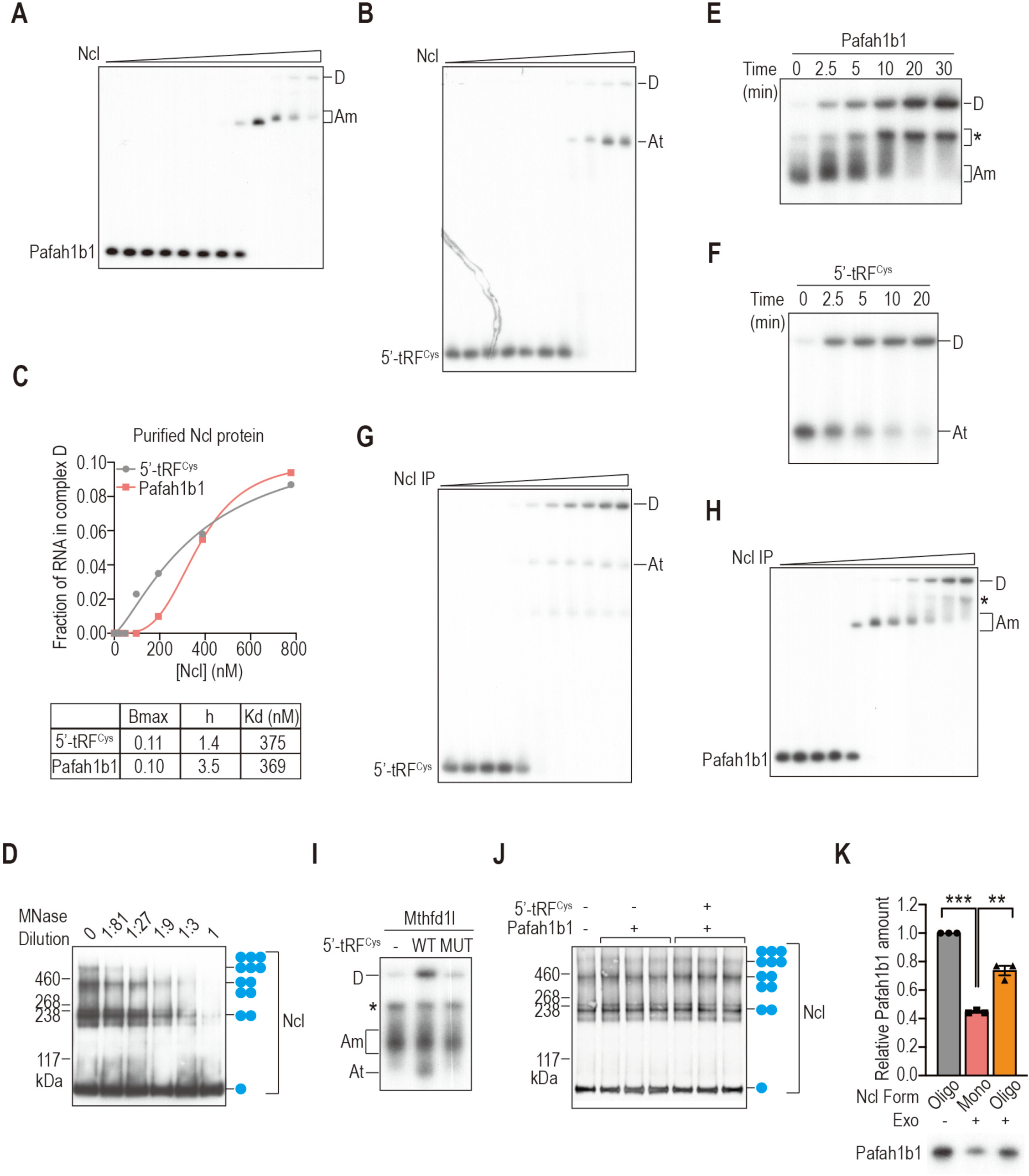
5’-tRF^Cys^ promotes complex D assembly and Nucleolin oligomerization. A, B. Native gel analysis of Nucleolin complexes assembled from Pafah1b1 (A) or 5’-tRF^Cys^ (B) using increasing amounts of Nucleolin protein. C. Quantification of complex D assembly as a function of Nucleolin concentration using purified Nucleolin protein. Bmax, specific maximum binding. h, Hill coefficient. Kd, equilibrium dissociation constant. D. Representative images of western blots of Nucleolin from Nucleolin IP that was pre-treated with different dilutions of micrococcal nuclease to remove endogenous RNAs before complexes were assembled at 30 °C and crosslinked with ethylene glycol bis (succinimidyl succinate). The number of blue dots represent the inferred number of Nucleolin monomers based on the molecular weight. E, F. Kinetics of Nucleolin complexes assembled from Pafah1b1 (E) or 5’-tRF^Cys^ (F) using Nucleolin IP. See also Figure 5F. Asterisk denotes an RNA-protein complex that was detected only with Nucleolin IP but not Nucleolin protein. G, H. Native gel analysis of Nucleolin complexes assembled from Pafah1b1 (G) or 5’-tRF^Cys^ (H) using increasing amount of Nucleolin IP. See also Figure 5G. I. Native gel analysis of Nucleolin complexes assembled using Nucleolin IP from Mthfd1l alone, or together with a wild-type (WT) or Nucleolin binding deficient (MUT) 5’-tRF^Cys^. Asterisk denotes an RNA-protein complex that was detected only with Nucleolin IP but not Nucleolin protein. J. Representative western blot of Nucleolin using Nucleolin IP incubated with or without Pafah1b1, or with both Pafah1b1 and 5’-tRF^Cy^ at 30 °C before crosslinking with EGS. See also Figure 5I. K. Top, quantification of the protection provided by different forms of Nucleolin from degradation by a prototypical 5’->3’ exonuclease Terminator after conducting the assembly assay at 4 °C or 30 °C to form monomeric Nucleolin (complex A) or oligomeric Nucleolin (complex D) respectively. Bottom, representative image of denaturing PAGE analysis of the exonuclease degradation products.

**Figure 6.**
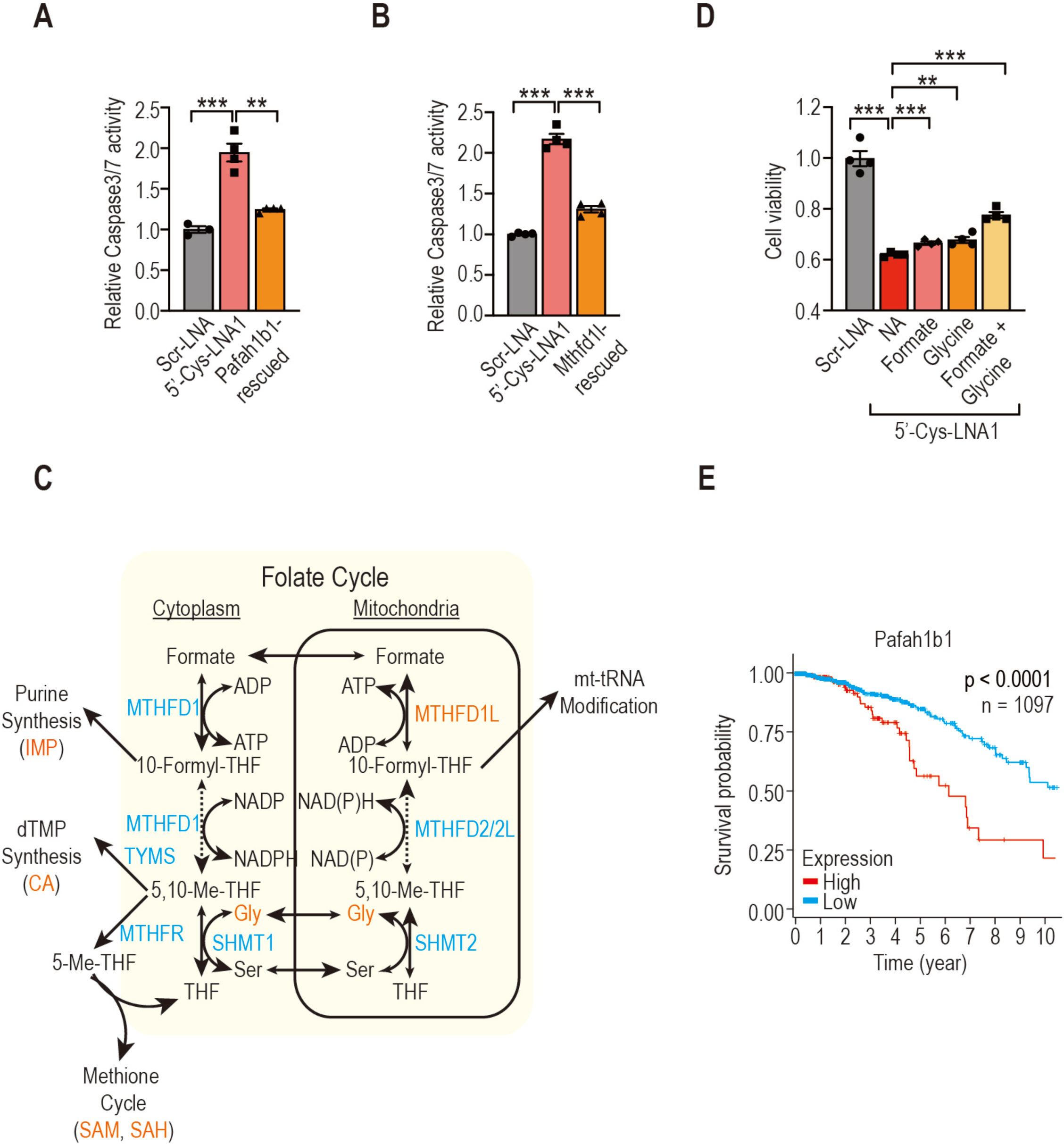
Pafah1b1 and Mthfd1l function downstream of 5’-tRF^Cys^ to promote breast cancer metastasis. A, B. Quantification of Caspase3/7 activity in 4T1 cells transfected with a control LNA (Scr-LNA), a 5’-tRF^Cys^ targeting LNA (5’-Cys-LNA1) either alone or together with overexpression of Pafah1b1 (Pafah1b1-rescued) (A) or Mthfdl1 (Mthfd1l-rescued) (B). Statistical significance was determined by one-tailed t-test with Welch’s correction. *, p<0.05; **, p<0.01; ***, p<0.001. C. Schematic overview of folate metabolism. Metabolic enzymes and key metabolites are indicated in blue and black, respectively. Metabolites that were detected by untargeted metabolite profiling to be upregulated in both the control and Mthfd1l-rescued cells are marked in orange (see also Figure 6C). Dashed lines represent two or more reactions. Arrowhead thickness denotes the preferred reaction direction. THF, tetrahydrofolate. Gly, glycine. Ser, serine. IMP, inosine monophosphate. CA, carbamoyl-aspartate. SAH, S-adenosylhomocysteine. SAM, S-adenosylmethionine. D. Relative viability of 4T1 cells two days post transfection with a control LNA (Scr-LNA), or a 5’-tRF^Cys^ targeting LNA (5’-Cys-LNA1) with or without supplementation of key metabolites in the folate cycle (1 mM formate and/or 2 mM glycine). Statistical significance was determined by one-tailed t-test with Welch’s correction. *, p<0.05; **, p<0.01; ***, p<0.001. E. Kaplan-Meier curves depicting survival probability of breast cancer patients in the TCGA cohort (n=1097) stratified by the expression of Pafah1b1. Statistical significance was determined by Mantel–Cox log-rank test.

